# Increasing Chemotherapeutic Efficacy Using pH Modulating and Doxorubicin Releasing Injectable Chitosan-Polyethylene Glycol Hydrogels

**DOI:** 10.1101/2023.07.06.547993

**Authors:** Zahra Ahmed, Kevin LoGiudice, Gavin Mays, Angelina Schorr, Rachel Rowey, Haisong Yang, Shruti Trivedi, Vikas Srivastava

**Author notes:** E-mail: vikas.

## Abstract

Modulation of pH is crucial to maintaining the chemical homeostasis of biological environments. The irregular metabolic pathways exhibited by cancer cells result in the production of acidic byproducts that are excreted and accumulate in the extracellular tumor microenvironment, reducing its pH. As a consequence of the lower pH in tumors, cancer cells increase the expression of metastatic phenotypes and chemotherapeutic resistance. A significant limitation in current cancer therapies is the inability to locally deliver the chemotherapy, leading to significant damage to healthy cells in systemic administration. To overcome these challenges, we present an injectable chitosan-polyethylene glycol hydrogel that is dual-loaded with doxorubicin and sodium bicarbonate providing alkaline buffering of extracellular acidity and simultaneous chemotherapeutic delivery to increase chemotherapeutic efficacy. We conducted in vitro studies of weak base chemotherapeutic and alkaline buffer release from the hydrogel. The release of doxorubicin from hydrogels increased in a low pH environment and was dependent on the encapsulated sodium bicarbonate concentration. We investigated the influence of pH on doxorubicin efficacy and viability of MCF-7 and MDA-MB-231 breast cancer cell lines. The results show a 2 to 3 fold increase in IC_50_ values from neutral pH to low pH, showing decreased cancer cell viability at neutral pH as compared to acidic pH. The IC_50_ results were shown to correlate with a decrease in intracellular uptake of doxorubicin at low pH. The proposed hydrogels were confirmed to be non-toxic to healthy MCF-10A mammary epithelial cells. Rheological studies were performed to verify that the dual loaded hydrogels were injectable. The mechanical and release properties of the hydrogels were maintained after extended storage. The chemotherapeutic activity of doxorubicin was evaluated in the presence of the proposed pH regulating hydrogels. The findings suggest a promising non-toxic, biodegradable hydrogel buffer delivery system that can achieve two simultaneous important goals of local acidosis neutralization and chemotherapeutic release.

## Introduction

Acidosis of the cancer microenvironment is a well-known characteristic of solid tumors.^1–3^ Tumor tissues contain an irregular network of blood vessels, causing spatially heterogeneous delivery of oxygen and nutrients to tumor cells.^4^ Loss of uniform oxygen perfusion and increased demand for cell division stimulates cancer metabolism to upregulate glycolytic pathways.^5–8^ To avoid inhibitory influences of intracellular acidification on metabolism and biological function, cancer cells equip themselves with well-developed defense systems of net acid extruders (e.g., Na^+^/HCO_3_ co-transporters, Na^+^/H^+^ exchangers, and H^+^-ATPases) and monocarboxylate transporters (e.g., MCT1 and MCT4) that extrude acid equivalents from the cytosol to the extracellular space.^9–11^ The abundance of these acid efflux transporters in cancer cells allows for a relatively neutral intracellular pH to be maintained despite the excessive acid producing metabolic processes.^12^ Greater diffusion distances between blood vessels and tumors prevent efficient venting of acid efflux. The body’s inherent bicarbonate pH buffering systems, which interact with acids to produce pH-neutral CO_2_ and H_2_O molecules, are overridden by lactic acid production.^13,14^ A consequence of acid accumulation is low extracellular pH (pH *≈* 6.5–6.8), which is recognized as a hallmark of cancer progression.^15,16^ The acidity of the tumor microenvironment produces a natural selection of cellular mutations favoring the most aggressive phenotypes.^17–19^ Cancer cells that proliferate in the low pH environment express increased invasion, metastasis, immune evasion, and chemoresistance.^20–22^ Cancer therapies, including chemotherapeutics, are negatively affected by the acidic extracellular pH through reduced drug uptake, increased drug efflux from cancer cells, or altered drug metabolism.^20^ In particular, it has been demonstrated that extracellular acidity reduces the efficacy of weak base chemotherapeutics, such as doxorubicin, mitoxantrone, paclitaxel, vinblastine, and vincristine.^23^ Ionization of the extracellular space renders weak base chemotherapeutics, a predominant class of cancer treatment drugs, less capable of permeating through cell membranes due to protonation of drug.^24–26^ This phenomenon is termed “ion trapping” and is responsible for the suppression of drug cytotoxicity, leading to a drugresistant cancer cell phenotype. Therefore, the acidity of the tumor microenvironment can be combated to enhance chemotherapeutic efficacy.

Various approaches have been made to prevent or negate the effects of extracellular acidity. The most direct approach has been achieved by neutralizing the pH through the administration of alkaline agents, such as bicarbonates or other bases.^5,27^ Studies in mice have shown that buffering of the tumor microenvironment through oral administration of bicarbonate has inhibited metastasis in breast cancer models.^28,29^ In a transgenic prostate cancer mouse model, administration of sodium bicarbonate at 4 weeks of age, significantly delayed the onset of cancers, while administration after 10 weeks inhibited the development of metastases.^30^ However, with oral delivery, only a very small proportion of sodium bicarbonate can reach the tumor at biologically tolerable oral dosages.^31^ For translational application in humans, the equivalent daily dosage of oral sodium bicarbonate (based on mouse trials) was calculated to be 13 to 15 g for a 70 kg human.^32^ Although feasible, in clinical trials, patient compliance of oral administration of sodium bicarbonate is relatively low because of gastrointestinal tract irritability, poor taste, or excessive capsule ingestion.^33,34^ The necessity for continuous administration of a buffering agent over time to maintain therapeutic efficacy while avoiding alkalosis poses a barrier to the implementation of this adjuvant therapy. Intravenous delivery of sodium bicarbonate-loaded liposomes and doxorubicin liposomes in a mouse model with xenogenic 4TI aggregates was shown to reduce tumor size and inhibit lung metastasis.^35^ In addition, calcium carbonate nanoparticles have been shown to decrease the proliferation and invasion of breast cancer cells in vitro.^36^ However, systemic delivery methods remain challenging due to the potential premature release of sodium bicarbonate in circulation and poor tumor specificity that increase the risk of alkalinity at off target sites. Similarly, intravenous delivery methods of doxorubicin can induce off-target toxicity in healthy cells due to inadequate tumor penetration.^37–41^ Sodium bicarbonate + doxorubicin microspheres and calcium carbonate + doxorubicin microspheres have been utilized to improve chemoembolization treatment in hepatocellular carcinoma.^42,43^ This method was developed specifically for liver cancer treatment by incorporating sodium bicarbonate and chemotherapeutics delivery through an arterial catheter.

There is a need for biocompatible, biodegradable, injectable, and local delivery methods combining chemotherapeutic delivery with pH regulation to increase chemotherapeutic efficacy and overcome the drawbacks of current systemic delivery methods. *For local delivery of the required amount of buffer to maintain physiological extracellular pH, we applied an injectable semi-interpenetrating network of chitosan and polyethylene glycol (PEG) hydrogel delivery system that can achieve efficient buffering of the cancer microenvironment through local release of sodium bicarbonate, an alkaline agent, and doxorubicin, a drug, for chemotherapeutic applications (Scheme 1*). Hydrogels were fabricated by covalent cross-linking of chitosan using genipin and physical cross-linking of chitosan with sodium bicarbonate. Chitosan was selected due to its biocompatibility, biodegradability, and its pH sensitivity. ^44,45^ PEG was incorporated into the hydrogel to enhance the mechanical stability and control the hydrogel degradation rate.^46^ The hydrogel cross-linking, rheological properties and surface morphology were investigated to understand and characterize the effects of physical and covalent dual cross-linking mechanisms. Swelling along with release studies helped understand pH dependent release of doxorubicin and sodium bicarbonate from the formulated chitosan-PEG hydrogels. Rheological investigations were carried out to assess hydrogel injectability. The release and rheological properties were also evaluated after an extended storage period of the hydrogel components to validate preservation of mechanical properties and drug potency. The cytotoxicity and cytocompatibility of unloaded and bicarbonate-loaded hydrogels were assessed using a mammary non-tumorogenic epithelial cell line (MCF-10A). Cell viability and drug uptake were assessed in MDA-MB-231 and MCF-7 cancer cells using MTT assay and confocal microscopy, respectively.

**Scheme 1:**
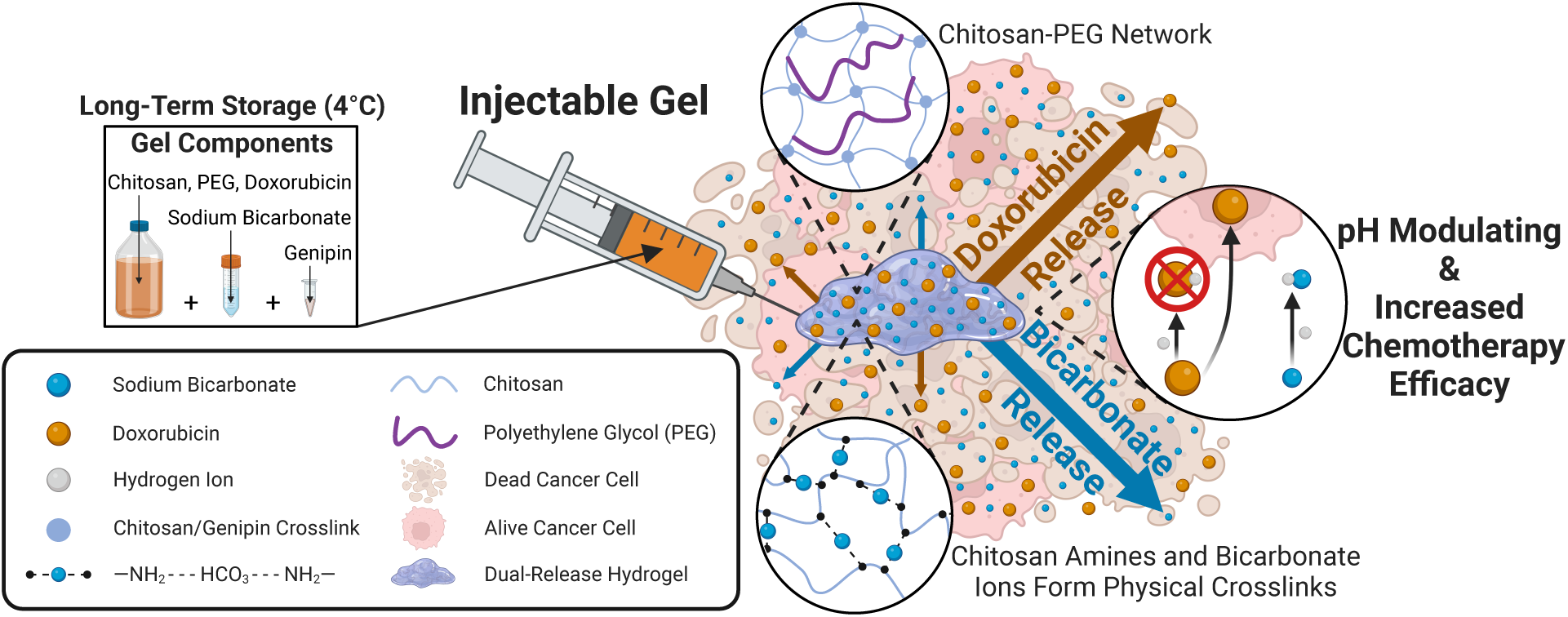
Injectable chitosan-PEG hydrogel for dual delivery of sodium bicarbonate and doxrubicin for increasing chemotherapy efficacy.

## Results and Discussion

### Rheological and morphological properties of chitosan-PEG hydrogels

We developed a biocompatible, biodegradable, semi-interpenetrating network of chitosan and PEG hydrogel to increase chemotherapeutic efficacy by providing local delivery of sodium bicarbonate and doxorubicin. Sodium bicarbonate was selected as a biocompatible molecule to increase pH of the acidic environment. The hydrogels were covalently cross-linked with genipin. Including sodium bicarbonate during the gel fabrication process added a secondary benefit of physical cross-linking in addition to the primary mechanism of covalent cross-linking (Fig. 1A). The covalent cross-links result from a nucleophilic substitution reaction mechanism between the amino group of chitosan and the olefinic carbon atom of genipin, forming a heterocyclic amine.^47^ The subsequent reaction is the formation of an amide through the reaction of the amino group on chitosan with the ester group of genipin.^47^ Sodium bicarbonate has been previously shown to react with cationic amino groups on chitosan.^48,49^ Neutralization promotes electrostatic repulsion among chitosan molecules, allowing hydrogen bonds and hydrophobic interactions to form between chitosan chains. ^50,51^ The covalent polymerization interaction between genipin and chitosan produces dark blue coloration, this was visibly reduced for hydrogels with increasing sodium bicarbonate concentrations (Fig. 1A), indicating the formation of hydrogels with both covalent and physical cross-linking networks. The hydrogels were designated as −DOX to indicate no doxorubicin or +DOX to indicate the presence of 50 µM doxorubicin in the experiments. The numbers 0, 50, 100, 150, and 200 following DOX indicate the concentration of sodium bicarbonate in mM. The concentration of doxorubicin (50 µM) loaded into 100 mm^3^ volume hydrogels was selected based on cell viability and release experiments. This loading allows for a sufficient released concentration of doxorubicin in release media to produce a therapeutic effect.

**Figure 1:**
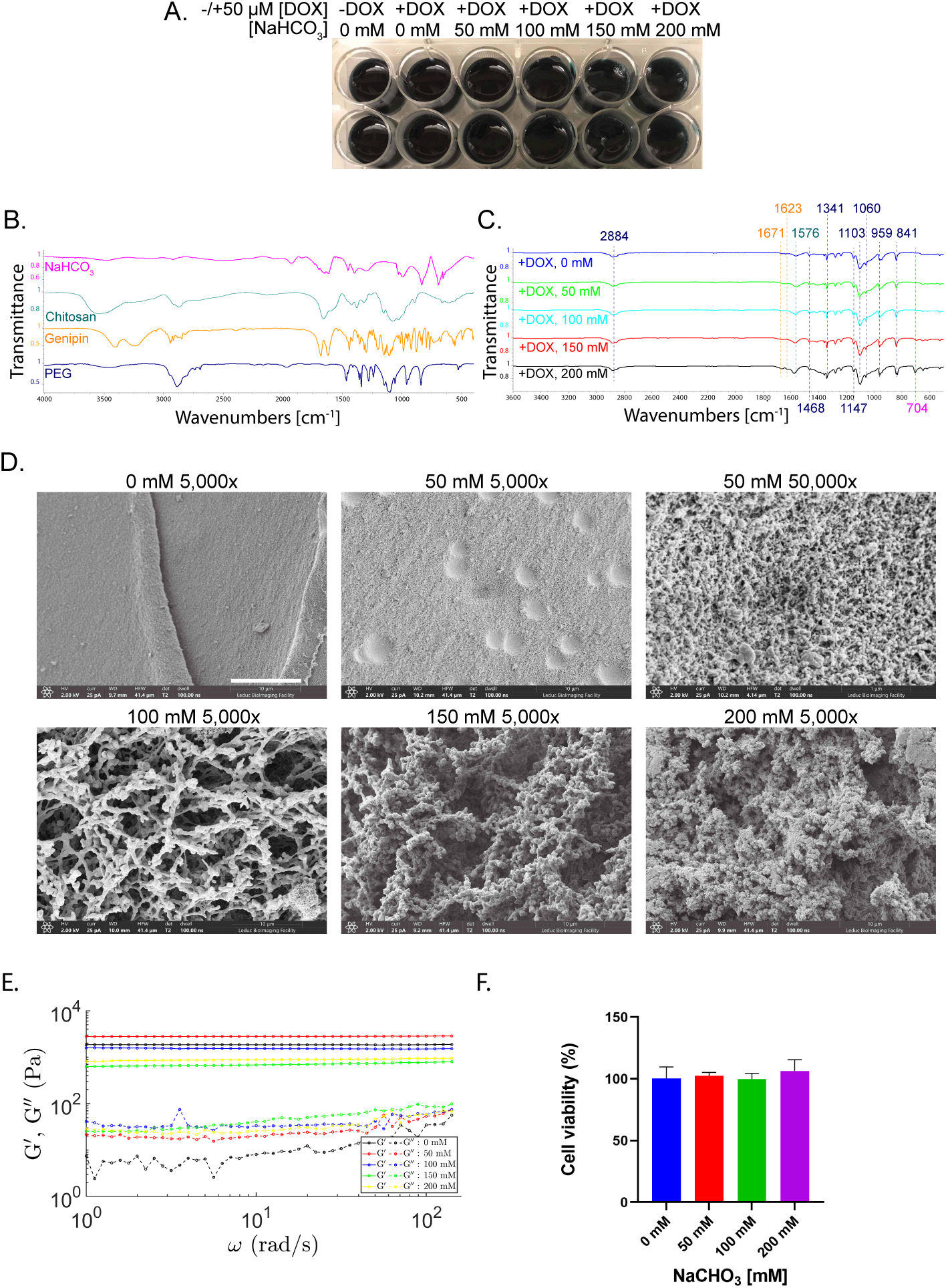
Characterization of doxorubicin/sodium bicarbonate loaded chitosan-PEG hydrogels formed at 37°C for 24 h. (A) Macroscopic view of chitosan-PEG hydrogels. The hydrogels were designated as −DOX to indicate no doxorubicin or +DOX to indicate the presence of 50 µM doxorubicin. The numbers 0, 50, 100, 150, and 200 following DOX indicate the concentration in mM of sodium bicarbonate present in 100 mm^3^ volume hydrogels. (B) FTIR spectra of individual hydrogel constituents. (C) FTIR spectra of hydrogel samples showing bonds after cross-linking. (D) SEM of the cross-sections of doxorubicin loaded hydrogels with varying sodium bicarbonate concentrations (0-200 mM) showing morphology at magnification 5,000x (50,000x included for pore visualization of 50 mM case). Scale bar: 10 µm and 1 µm for 5,000x and 50,000x, respectively. (E) Oscillatory shear experiment at 1% strain. (F) Toxicity of NaHCO_3_-loaded chitosan-PEG hydrogels quantified using MTT viability assay on MCF-10A cells after 24 h of exposure. p *>* 0.05, for all cases. All values are expressed as mean ± SE (n=6).

FTIR spectra were recorded to determine the structural changes in the cross-linked hydrogel upon the interaction between chitosan, PEG, genipin and sodium bicarbonate. The characteristic bands for the individual constituents (sodium bicarbonate, chitosan, genipin, and PEG) are presented in Fig. 1B. The resulting spectra for the hydrogels after 24 h of cross-linking (+DOX, 0 mM, +DOX, 50 mM, +DOX, 100 mM, +DOX, 150 mM, +DOX, 200 mM) is shown in Fig. 1C. The recorded hydrogel spectra (Fig. 1C) shows that characteristic peaks of both chitosan and PEG are observed in the hydrogel spectra, confirming that cross-linking the chitosan in the presence of PEG resulted in a semi interpenetrating network hydrogel. The spectrum of uncross-linked chitosan shows absorption’s at 1660 cm*^−^*^1^ and 1595 cm*^−^*^1^, characteristics of C-O stretching and N-H bending vibrations of the amide II present on chitosan’s molecular structure.^47,52,53^ After chitosan cross-linking with genipin, the amide II band shifted to 1576 cm*^−^*^1^ due to the formation of secondary amide linkages. The FTIR spectrum of genipin (Fig. 1B) showed two strong bands at 1671 cm*^−^*^1^ which correspond to C=C stretching of the cyclic alkenes and a peak at 1623 cm*^−^*^1^ representing C-C stretching of the cycloolefin.^54^ These peaks intensify for +DOX, 150 mM NaHCO_3_ and +DOX, 200 mM NaHCO_3_ hydrogel samples, suggesting an increase in the amount of uncross-linked genipin with increasing sodium bicarbonate concentrations. The spectrum of PEG showed a peak at 2888 cm*^−^*^1^ assigned to C-H stretching, 1468 cm*^−^*^1^ and 1342 cm*^−^*^1^ assigned to C-H bending, 1104 cm*^−^*^1^ as a characteristic of C-O-H stretching, and 962 cm*^−^*^1^ and 842 cm*^−^*^1^ attributed to C-C chain vibration.^55^ These peaks are all retained in cross-linked hydrogels.

The morphologies of the synthesized hydrogels were obtained by scanning electron microscope (SEM) images of the cross sections of critical-point-dried hydrogels that were loaded with doxorubicin at increasing sodium bicarbonate concentrations. As seen in (Fig. 1D), the pore size of the hydrogels increased with increasing concentrations of sodium bicarbonate. No visible pores were seen for hydrogels without sodium bicarbonate at 5,000x or 50,000x magnification. As shown previously by Vo et. al., the addition of PEG to genipin cross-linked hydrogels decreases the pore size significantly. ^55^ At low concentrations of sodium bicarbonate (50 mM), pores of *≤* 50 nm diameter can be seen at 50,000x magnification. With increasing concentrations of bicarbonate, the pore size significantly increases to µm length scales.

To examine the effect of the sodium bicarbonate content on the mechanical properties of chitosan/doxorubicin hydrogel systems, we performed oscillatory shear experiments on fully cross-linked hydrogels to obtain the storage modulus (G’), representing the elastic behavior of the material, and loss modulus (G”), indicative of the viscous behavior (Fig. 1E). The storage modulus and loss modulus of the hydrogels were measured using a rheometer with a parallel plate geometry at the angular frequency range of 1 to 100 rad/sec. The G’/G” ratio, of 30 was maintained over this frequency range, indicating more elastic solid-like behavior and frequency independence. Frequency independence is maintained for all bicarbonate concentrations. The addition of sodium bicarbonate from 50 to 200 mM decreases the storage modulus of the hydrogels. This can be attributed to sodium bicarbonate’s gelation mechanism with chitosan. Chitosan forms physical junctions as a result of sodium bicarbonate neutralization of chitosan amine groups. Gelling systems containing sodium bicarbonate produce a combination of covalent and physical entanglements.

To validate the use of the chitosan-PEG hydrogels for a potential tumor delivery system, we assessed the cytotoxicity to non-tumorigenic mammary epithelial cells using an MTT assay (Fig. 1F). The MCF-10A breast cells were exposed to chitosan-PEG hydrogels loaded with 0, 50, 100, and 200 mM sodium bicarbonate (no doxorubicin) over 24 h. The hydrogels showed good cytocompatibility as there were no significant differences in cell viability when normalized to the control case without hydrogel (*≥* 100% cell viability). There was also no significant differences across all hydrogel cases, indicating no toxicity from sodium bicarbonate (Fig. 1F).

### Injectability of chitosan-PEG hydrogels

In contrast to systemic administration, localized delivery of sodium bicarbonate and doxorubicin enhances drug distribution while minimizing toxicity to normal tissues. To enable this potential localized delivery, we evaluated the injectability of the chitosan-PEG hydrogels. The covalent cross-linking mechanism of genipin with chitosan occurs slowly over the duration of 24 h.^46^ The vial tilting method demonstrated that genipin-crosslinked hydrogels, without sodium bicarbonate, retained their flowability even after 30 minutes of crosslinking (Fig. 2A). A comparison time sweep of storage modulus of chitosan (1.5% w/v) hydrogel constituents without cross-linker and chitosan hydrogel cross-linked with genipin indicate that genipin only starts to initiate covalent cross-linking in hydrogels at 40 min of test duration (storage modulus of chitosan with genipin and chitosan without are equivalent until 40 min). The addition of sodium bicarbonate to the hydrogels initiated physical cross-linking at shorter time scales (Fig. 2A) allowing for more rapid hydrogel formation after injection of liquid constituents, preventing a loss of material before covalent cross-linking occurs. Comparing tan*δ*, the ratio of the loss modulus over the storage modulus (G”/G”), the ability of a material to dissipate energy (viscous flow as tan*δ* increases) or to store energy (elastic solid as tan*δ* decreases). At the start of the test (t = 0), the hydrogel formulations have a higher tan*δ* (.2 to .5) which starts to plateau to smaller values at t = 400 s (.07 to .1) (Fig. 2B). Oscillatory rheological testing demonstrated that all the formulations exhibit a frequency independent storage modulus at 30 min of physical cross-linking (Fig. 2C). All hydrogel formulations with sodium bicarbonate experience a decrease in viscosity upon application of higher shear rate as more physical cross-links slide at lower rates leading and to a more viscous response (Fig. 2D). This also enables injection of shear-thinning hydrogels after physical cross-linking starts to occur. The viscosity values at higher injectible flow rates are in a flowable/injectable regime.

**Figure 2:**
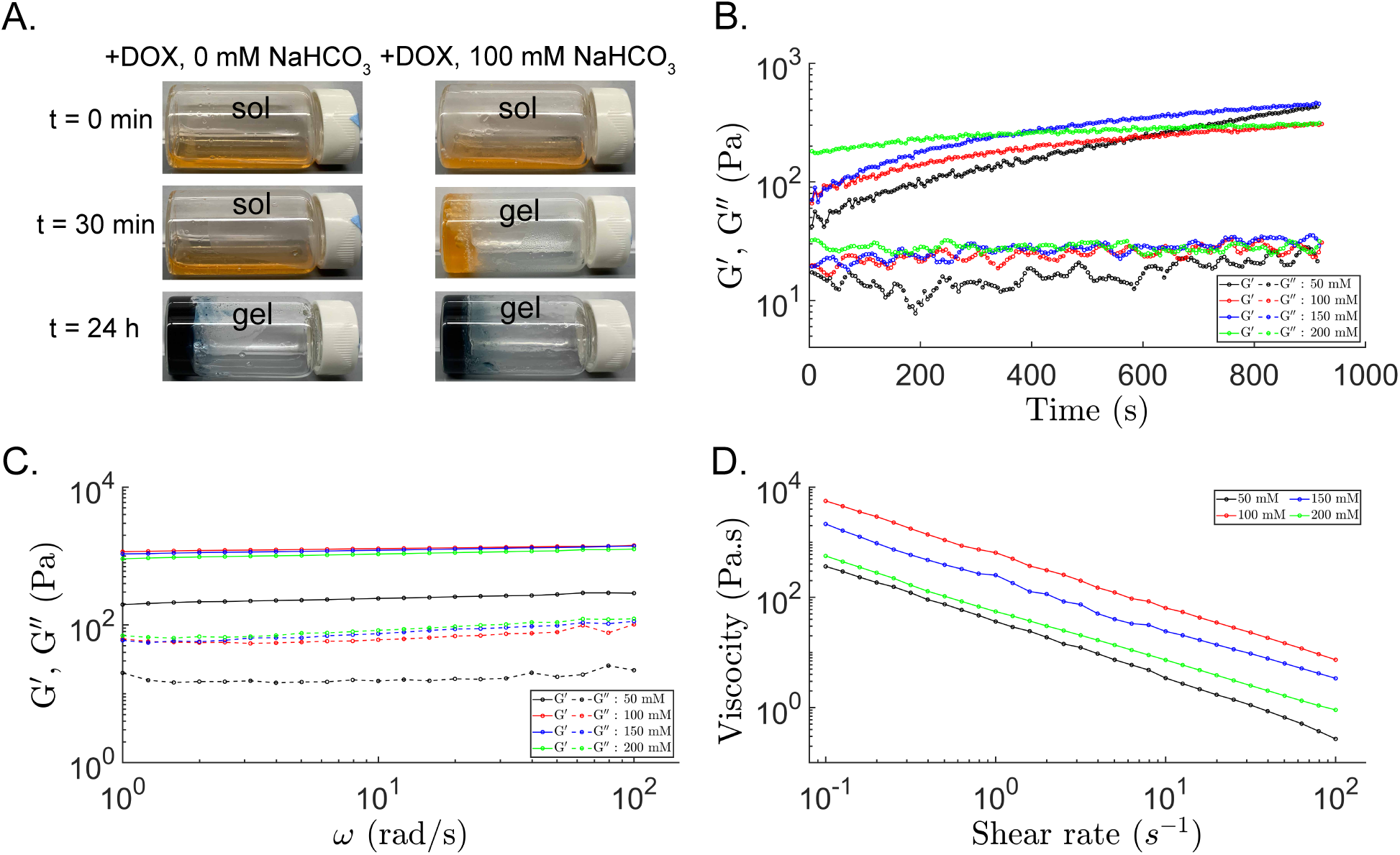
Rheological properties of chitosan-PEG hydrogels with sodium bicarbonate. (A) Hydrogel cross-linking with and without sodium bicarbonate visualized using the vial tilting method. (B) Time dependent response at 100 rad/s during hydrogel cross-linking. (C) Frequency sweeps at 1 % strain after 30 min of cross-linking. (D) Flow sweep from 0.1 to 100 (*s^−^*^1^) indicating shear-thinning properties after 30 min of cross-linking.

### Sodium bicarbonate and doxorubicin release from dual loaded chitosan-PEG hydrogels

After confirming hydrogel biocompatibility, we conducted sodium bicarbonate and doxorubicin release studies on dual loaded sodium bicarbonate and doxorubicin chitosan-PEG hydrogels. We obtained pH release profiles of the chitosan-PEG hydrogels loaded with varying concentrations of sodium bicarbonate. The starting solution was phosphate buffered saline (PBS) adjusted to pH 6 and then the loaded hydrogels were added. The representative pH of the acidic tumor microenvironment was considered to be 6.0.^15^ The control groups of hydrogels without sodium bicarbonate loading (+DOX, 0 mM NaHCO_3_) were used to confirm that there is no pH change produced by the hydrogel or doxorubicin alone. NaHCO_3_-loaded hydrogels caused a rapid increase in the pH of the solution within 8 h regardless of bicarbonate concentration (Fig. 3A). Following the initial burst release of the buffer from the hydrogel, the pH then gradually increased up to 120 h. The unloaded sodium bicarbonate hydrogel control group did not exhibit an increase in pH; a negligible decrease in pH is shown within the first 8 h due to the slightly acidic nature of the control hydrogel.

**Figure 3:**
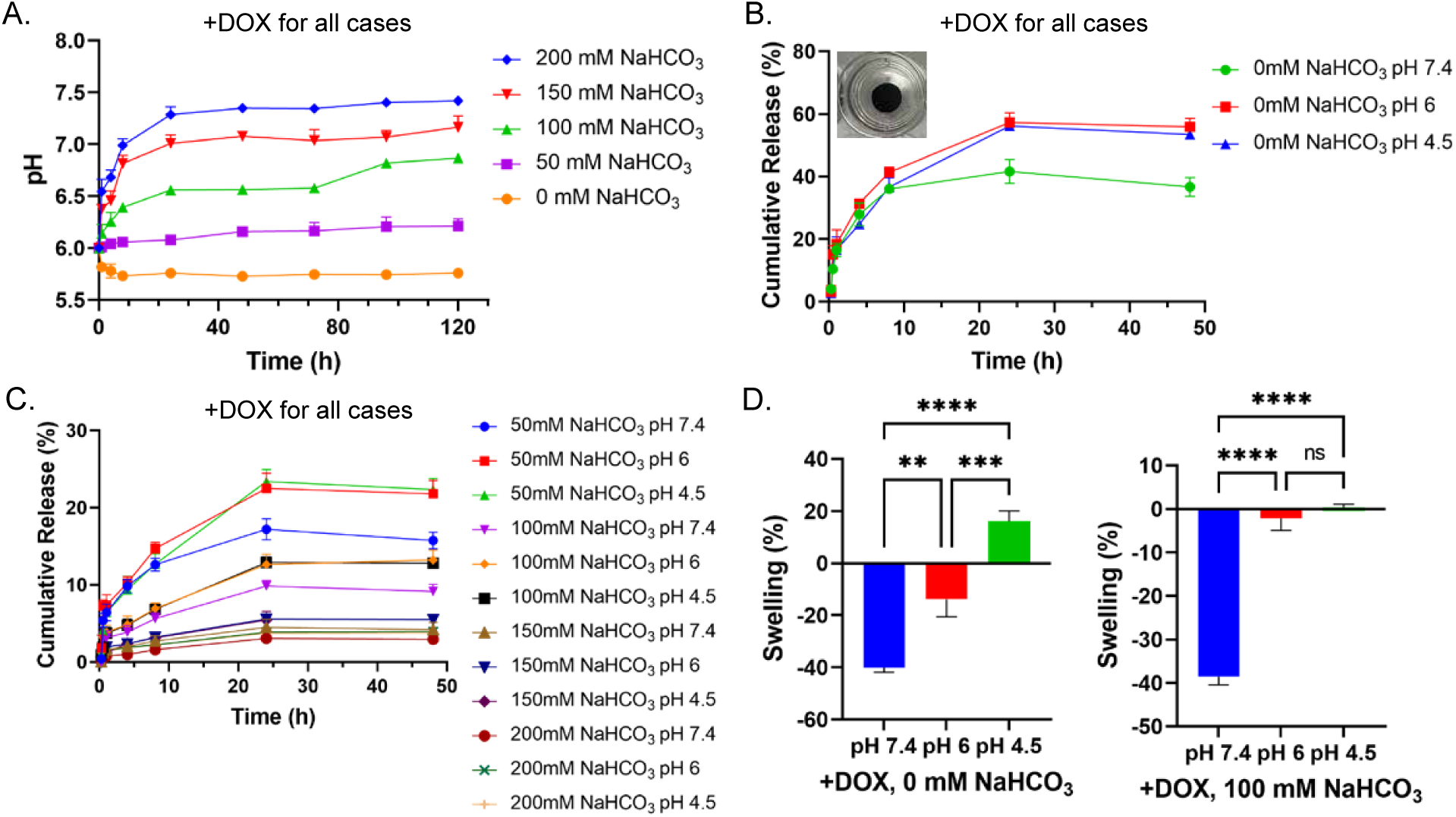
Sodium bicarbonate entrapment in hydrogels impacts pH and doxorubicin release. (A) Change in pH of PBS solution at starting pH 6 in the presence of doxorubicin (50 µM) and sodium bicarbonate (0-200 mM) loaded chitosan-PEG hydrogels. Release profile of chitosan-PEG hydrogels with (B) doxorubicin (50 µM) and (C) doxorubicin (50 µM) and sodium bicarbonate (50-200 mM) in PBS (pH 7.4, 6.0 and 4.5) after 48 h. (D) Swelling of doxorubicin chitosan-PEG hydrogels in PBS (pH 7.4, 6.0 and 4.5) after 24 h. Each value represents the mean ± SE (n=6) with significance defined as * p *<* 0.5; *** p *<* 0.001, **** p *<* 0.0001.

As our focus is on chemotherapeutic applications, a study was conducted to evaluate the in vitro release kinetic profiles of 50 µM of doxorubicin from 100 mm^3^ volume chitosan-PEG hydrogels. The loading concentration for doxorubicin was established such that it is measurably effective in our in vitro experiments on cancer cells. In our method, the doxorubicin dosage concentration can be adjusted within the hydrogel system. The pH values of 7.4, 6.0 and 4.5 were selected to mimic extracellular normal physiological conditions, tumor microenvironment conditions, and overly acidic conditions, respectively. The release of doxorubicin from the hydrogels was monitored by fluorescence reading at 37°C in 2 mL of release media. The concentration in the media was measured at selected time points up to 48 h after the addition of the hydrogel. The results in (Fig. 3B) reveal that doxorubicin release exhibited an initial burst release over the first 2 h followed by a slower release rate up to 24 h. A comparison of the results under different pH conditions reveals the pH sensitivity of drug release. Doxorubicin release was slowest under the neutral (pH 7.4) environments, with around 39% doxorubicin release from gels without sodium bicarbonate over a period of 24 h. Under both lower pH conditions, approximately 60% of the doxorubicin was released within 24 h. This observation indicates that the release of doxorubicin from chitosan hydrogels is acid responsive.

We also studied the effect of sodium bicarbonate loading on doxorubicin release rates in dual loaded hydrogels. The incorporation of sodium bicarbonate prevented the initial burst release of doxorubicin and instead allowed sustained release over 24 h (Fig. 3C). This has the benefit of preventing local doxorubicin toxicity of healthy tissues. Loading 50 mM of sodium bicarbonate into hydrogels decreased the release of doxorubicin to 16% at pH 7.4 and 22% for pH 6 in 24 h. Similarly, the 100 mM released 10% at pH 7.4 and 12% at pH 6.0. It is expected that the hydrogels will continue to exhibit slow release after 24 h. Additionally, for each bicarbonate concentration, the hydrogels at the lower pH solutions demonstrate a slightly higher percent release of doxorubicin, which is more ideal for applications in the acidic cancer microenvironment.

Next we examined the swelling properties of the doxorubicin-loaded hydrogels with either 0 mM or 100 mM sodium bicarbonate by placing the hydrogel samples in PBS solution (pH 4.5, 6, 7.4) for 24 h (Fig. 3D). Hydrogel masses were measured before and after swelling. The highest swelling 16.0% for 0 mM sodium bicarbonate was observed for hydrogels submerged in pH 4.5, while a significant shrinking of 40.1% occurred for hydrogels at pH 7.4. The hydrogel containing 100 mM of sodium bicarbonate experiences little swelling at pH 4.5 and significant shrinking at pH 7.4. This can be attributed to the release of sodium bicarbonate, causing an increase in the pH of the hydrogel environment. This trend suggests a swelling dependent release of doxorubicin. Sodium bicarbonate loading increases internal hydrogel pH, and the release of sodium bicarbonate along with doxorubicin increases external solution pH causing decreased swelling or shrinking of hydrogels. These results demonstrated the potential to achieve both pH regulation and pH-sensitive chemotherapeutic release from these hydrogel delivery systems.

### Shelf-storage of chitosan-PEG hydrogels

To enable long-term storage followed by the utilization of the injectable chitosan-PEG hydrogel solutions for therapeutic administration, we evaluated their ability to maintain physical properties and doxorubicin potency even after extended storage durations. The primary storage method entails keeping the doxorubicin and hydrogel constituents separately from the cross-linkers genipin and sodium bicarbonate as separate solutions within a refrigerated environment at 4 °C. Combining the cross-linkers with the hydrogel constituents for storage at 4 °C is not recommended as it will lead to partial (covalent) hydrogel cross-linking and reduce injectability of the hydrogel. During application, these solutions are mixed at room temperature immediately prior to injection of the hydrogel. Utilizing a dual-chamber syringe for storing the components separately for subsequent mixing at room temperature upon injection is a potential storage method. After 24 h cross-linking, the chitosan-PEG hydrogels consistently maintained their rheological properties even after 14 days of storage of hydrogel constituents (Fig. 4A). Our investigations also confirmed that the release of doxorubicin from the hydrogels remained uncompromised (Fig. 4C & 2D), allowing for the storage of injection solutions without potency loss for a duration of 14 days. We expect that the effective storage life for the hydrogel system is much longer than 14 days. As a secondary option, which is more energy-consuming, we also investigated combining all hydrogel constituents and subsequently freezing the mixture for storage at −20 °C. The frozen and stored samples were thawed, and the results indicated that the mechanical and doxorubicin release properties of the thawed and cross-linked freeze-stored samples were identical to those of the freshly prepared and tested hydrogels.

**Figure 4:**
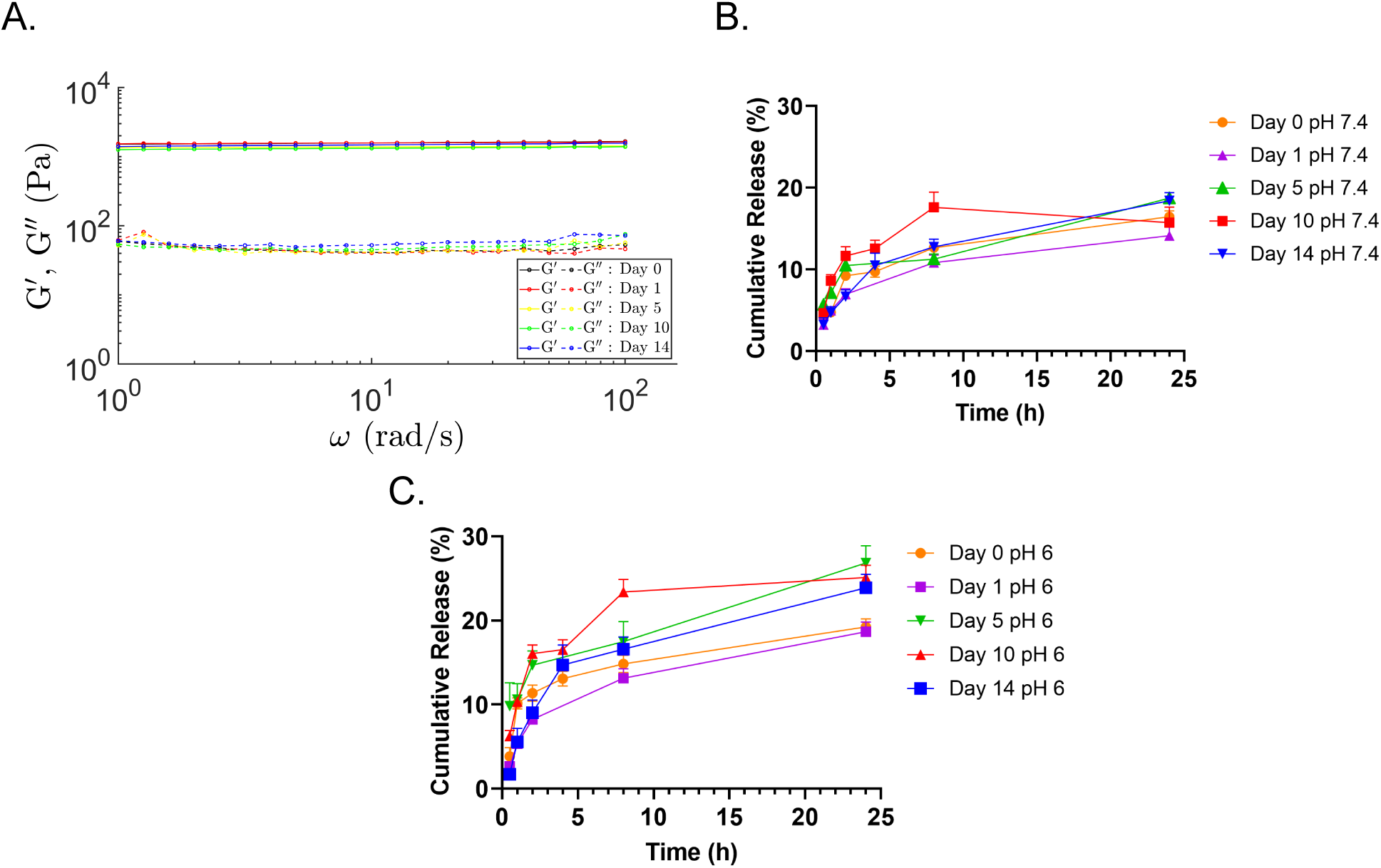
Rheological properties and doxorubicin release of cross-linked +DOX, 100 mM NaHCO_3_ chitosan-PEG hydrogels after 14 days storage of hydrogel constituents before cross-linking. (A) Frequency sweep at 1% strain. Release of doxorubicin after storage at 4 °C at pH (B) 7.4 and (C) 6. The mechanical properties and doxorubicin release are maintained after 14 days of storage of hydrogel constituents at 4 °C.

### pH-dependent cell viability and cell uptake

Ion trapping of weak base chemotherapeutics depends on the extracellular acidity in the tumor microenvironment.^24,25^ At first, we demonstrated the effect of pH on chemotherapeutic efficacy of doxorubicin by exposing individual cultures of MDA-MB-231 and MCF-7 breast cancer cells to varying concentrations of doxorubicin (0-10 µM) at pH values corresponding to the acidic pH (6.5) of the tumor microenvironment and to the neutral physiological pH (7.4) of healthy breast tissue. The low pH media was prepared by titrating with lactic acid.^56^ Cells were allowed to adhere as a monolayer to the surfaces of a well-plate overnight in complete media before doxorubicin was added. After 24 h of treatment, doxorubicin was significantly more effective at decreasing cell viability at pH 7.4 compared to acidic pH. The results show a 2 to 3 fold increase in IC_50_ values at pH 6.5 in comparison to pH 7.4 for both cell lines (Fig. 5). The IC_50_ increased from 1.5 µM at pH 7.4 to 3.4 µM at pH 6.5 for MDA-MB-231 cells. Similarly, in MCF-7, the IC_50_ increased from 0.5 µM at pH 7.4 to 1.6 µM at pH 6.5. The IC_50_ results highlight a more pronounced pH-dependent effect on doxorubicin efficacy within the MDA-MB-231 cell line compared to the MCF-7 cell line. The diminished effectiveness of doxorubicin under acidic pH conditions has primarily been attributed to ion trapping. ^57^ However, a direct cellular consequence of acidic pH is associated with an elevated expression of drug resistance proteins and metastatic biomarkers, both of which have been linked to doxorubicin resistance.^22,58–64^ MDA-MB-231 cells exhibit high invasiveness and metastatic potential while MCF-7 cells show non-invasiveness. It is expected that the positive effects of acidic pH neutralization is more significant in cell lines that exhibit high metastatic potential and chemotherapeutic resistance. Consequently, MDA-MB-231 shows increased doxorubicin efficacy in the presence of pH regulation.

**Figure 5:**
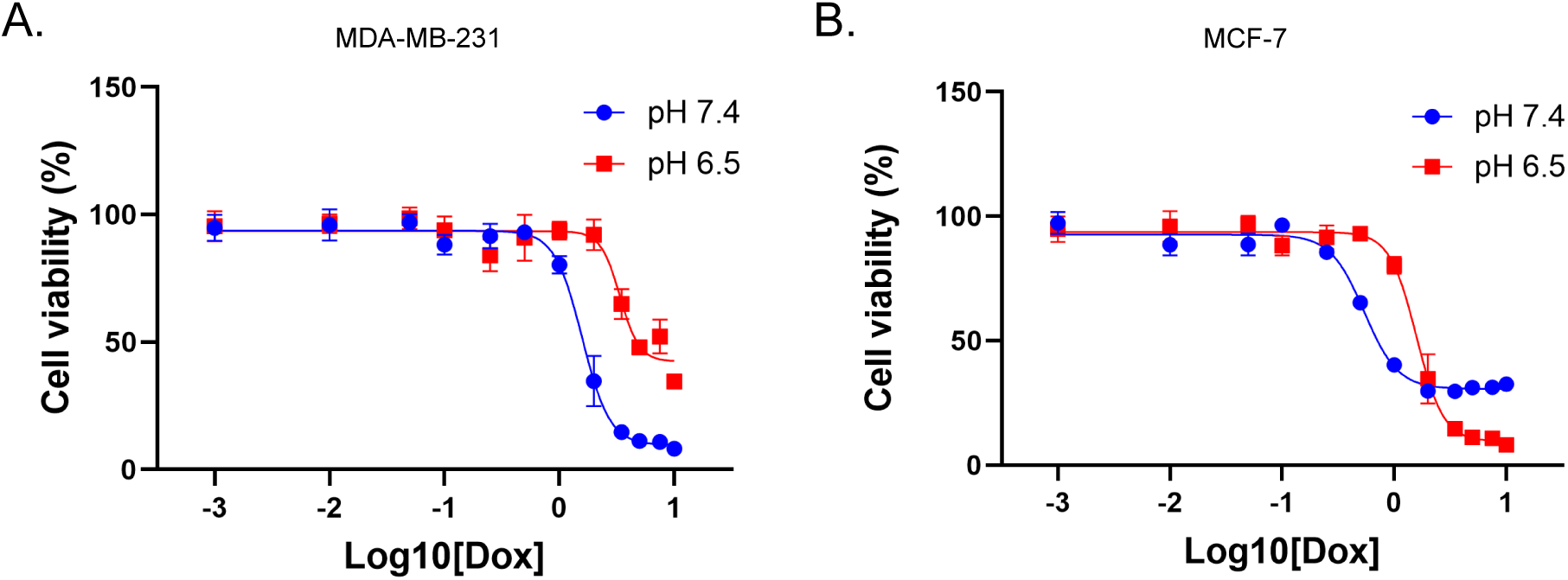
Cytotoxicity of doxorubicin is altered between physiological pH (7.4) and acidic pH (6.5). Varying concentrations (0-10 µM) of doxorubicin were administered, and MTT assays (n = 6) were conducted to study the effect on (A) MDA-MB-231 cells and (B) MCF-7 cells for 24 h. The IC_50_ values for MDA-MB-231 increase from 1.5 µM at pH 7.4 to 3.4 µM at pH 6.5. The IC_50_ values for MCF-7 are .50 µM at pH 7.4 and 1.6 µM at pH 6.5. Values for IC_50_ were calculated using GraphPad Prism.

After confirming the effects of pH on doxorubicin efficacy, we studied the hydrogel’s ability to prevent ion trapping and increase chemotherapeutic efficacy in acidic environments for controlled amounts of doxorubicin. Free doxorubicin and hydrogels loaded with or without 200 mM sodium bicarbonate were incubated with individual cultures of MDA-MB-231 and MCF-7 breast cancer cells at pH 6.5 media. The 200 mM sodium bicarbonate loaded hydrogel was selected to maintain a pH of 7.4 in its surrounding environment, as shown in Fig. 3A. After 48 h of treatment, we quantified the effect of incorporating NaHCO_3_-loaded hydrogels using MTT assay. Results showed that free doxorubicin, concurrent with sodium bicarbonate loading of hydrogels, significantly decreased cancer cell viability by reducing the effects of ion trapping compared to doxorubicin without sodium bicarbonate incorporation (Fig. 6). NaHCO_3_-loaded hydrogels reversed the effects of acidic pH on doxorubicin efficacy.

**Figure 6:**
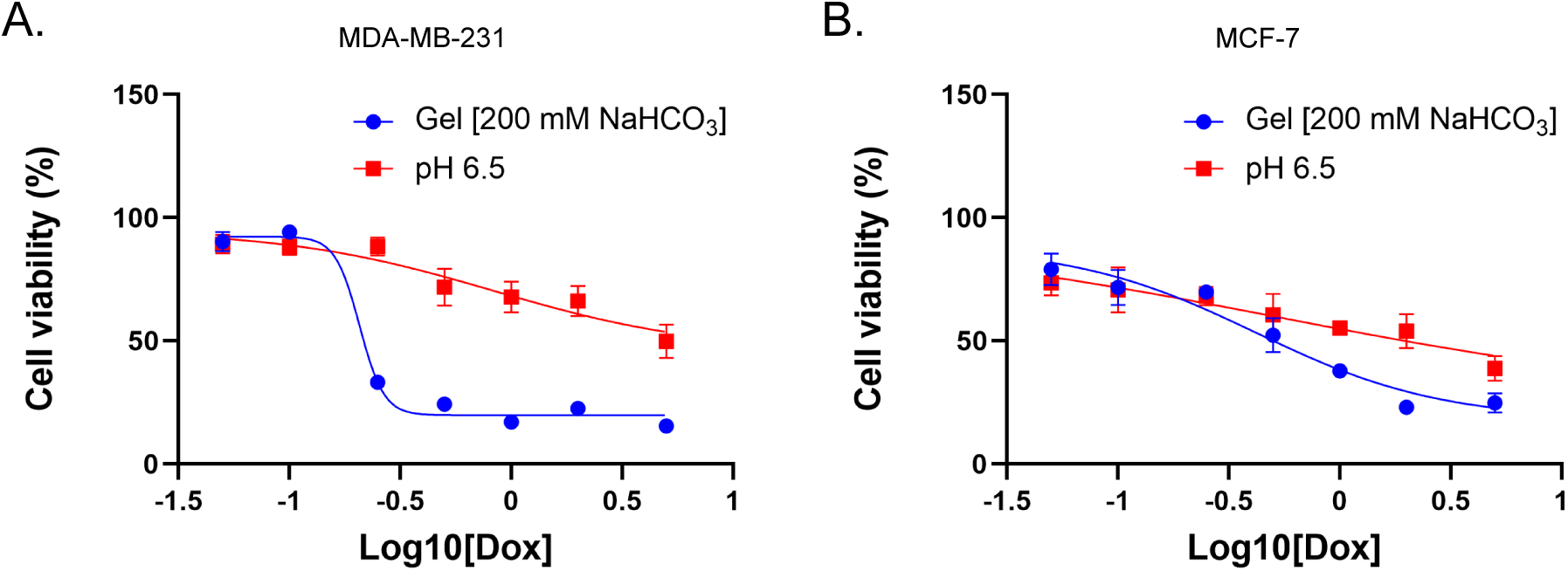
Comparison of cytotoxic effect at acidic pH (6.5) and pH achieved after addition of NaHCO_3_ loaded hydrogels (200 mM) to cultures of (A) MDA-MB-231 and (B) MCF-7 cell lines for 48 h. The IC_50_ values for MDA-MB-231 were calculated as 0.21 *µ*M with hydrogels and 0.84 µM at pH 6.5. The IC_50_ values for MCF-7 are 0.42 µM at pH 7.4 and 0.53 µM at pH 6.5.

An investigation was also made to determine the pH-dependent cellular internalization of doxorubicin. First, MDA-MB-231 cells and MCF-7 cells were individually cultured in fibronectin-coated glass bottom petri dishes in complete DMEM/F12 media. Then, adhered MDA-MB-231 and MCF-7 cells were co-incubated with hydrogels loaded with 200 mM of sodium bicarbonate in an acidic culture medium (pH 6.5) for 48 h. Free doxorubicin at the same pH of 6.5 was used as a comparison. Doxorubicin accumulation in cells was visualized by confocal fluorescence microscopy (Fig. 7A-B). The merged images for cases with free doxorubicin present showed that doxorubicin fluorescence signals were distributed in the nuclei and cytoplasm of MDA-MB-231 cells. While for MCF-7 cells, doxorubicin was primarly localized in the nucleus. MDA-MB-231 cells exhibit an increased uptake of doxorubicin into the cell cytoplasm. This accumulation within the cytoplasm has previously been associated with the development of doxorubicin resistance.^62,65–67^ For both cell lines, the intracellular accumulation of doxorubicin remained greater across all doxorubicin concentrations for hydrogel-induced high pH conditions in comparison to the low pH case. A quantitative analysis of fluorescence intensity in the cell nuclei (Fig. 7C-D) of MDA-MB-231 and MCF-7 cells reveals that the fluorescence intensity in the nuclei of the cells treated with free DOX at low pH was significantly lower than for cells treated with sodium bicarbonate loaded hydrogels. This revealed higher cellular accumulation of doxorubicin when it is present with sodium bicarbonate loaded hydrogels. The release of sodium bicarbonate from our gels increases the acidic pH to physiological pH and significantly reduces ion trapping of doxorubicin.

**Figure 7:**
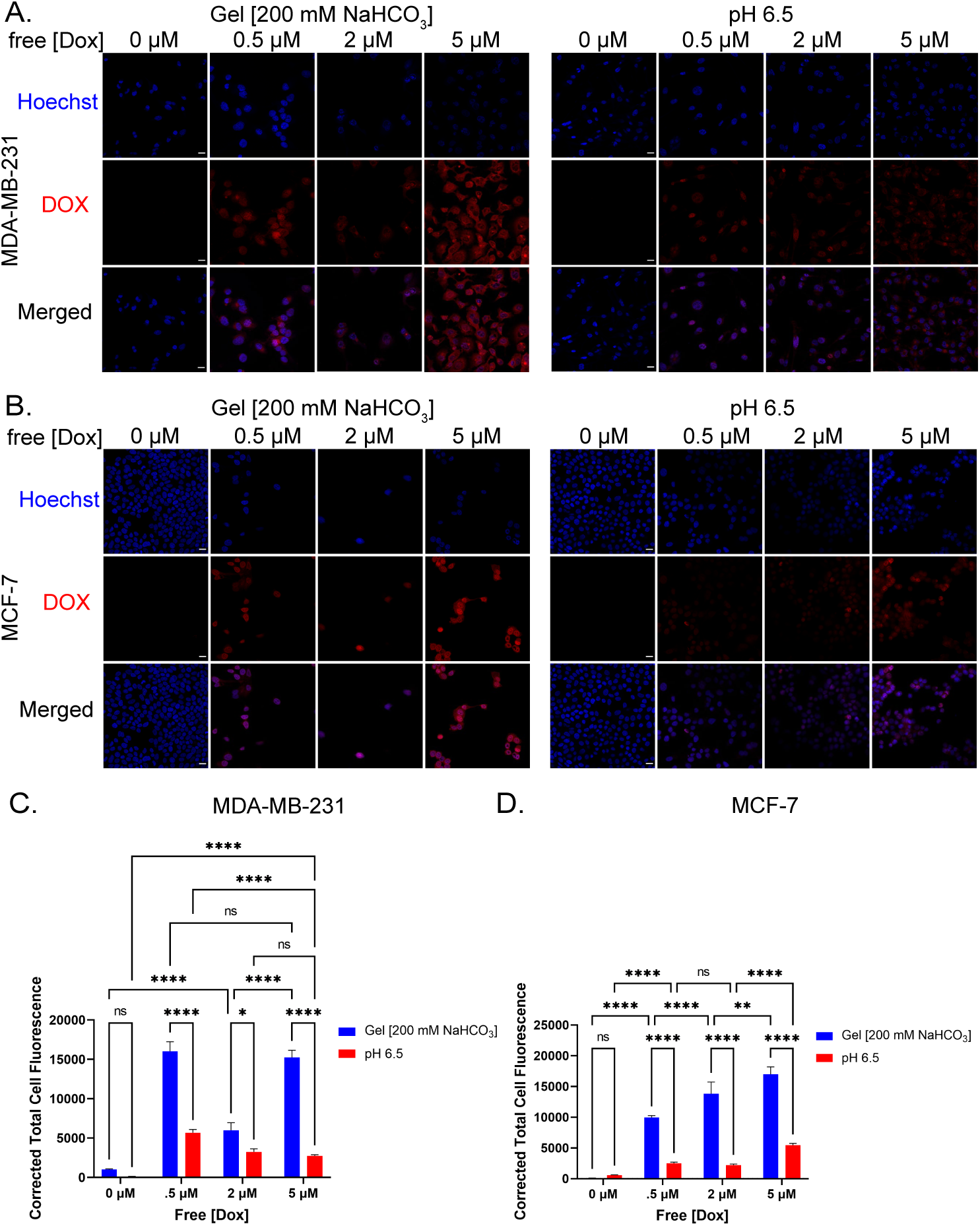
pH dependent localization of doxorubicin in the nucleus. Confocal fluorescence images of Hoechst stained (A) MDA-MB-231 and (B) MCF-7 cells treated with free doxorubicin (0-5 µM for 48 h). Left corresponds to treatment at high pH (induced by 200 mM sodium bicarbonate loaded hydrogels), and the right corresponds to cells treated at low pH (6.5) The red fluorescence of DOX was captured in the red channel, and Hoechst stained nuclei were captured in the blue channel. The overlapping images demonstrated the co-localization of DOX and nuclei, indicating the diffusion of DOX into nuclei and accumulation. Scale bar in left images: 20 µm. The quantitative analysis of fluorescence images of Hoechst stained (C) MDA-MB-231 and (D) MCF-7 cells treated with free doxorubicin (0-5 µM). The fluorescence intensity of DOX was calculated in the cell nucleus region via the red channel. All of the hydrogel treated cells demonstrated enhanced doxorubicin accumulation in cell nuclei at each concentration when compared to low pH (6.5). Each value represents the mean ± SE (n= 29-237, MDA-MB-231) and mean ± SE (n= 28-192, MCF-7) with significance defined as * p *<* 0.5; *** p *<* 0.001, **** p *<* 0.0001.

### Cellular uptake of doxorubicin from doxorubicin and bicarbonate loaded chitosan-PEG hydrogels

We evaluated the cellular internalization of doxorubicin released from dual loaded sodium bicarbonate and doxorubicin hydrogels (+DOX, 100 mM NaHCO_3_) using MDA-MB-231 and MCF-7 cell lines over 48 h (Fig. 8A-B). This hydrogel formulation was selected as it was shown to have optimal doxorubicin release while still allowing sufficient sodium bicarbonate release for pH regulation to near 7. The amount of doxorubicin released from the hydrogels was 0.47 *±* 0.05 µM after a 48 h in the culture. The +DOX, 100 mM NaHCO_3_ hydrogel was compared to free doxorubicin doses ranging from 0 to 5 µM at pH 6.5. Free doxorubicin was used as a control to establish the efficacy of the dual sodium bicarbonate and doxorubicin loaded hydrogel. The distribution of doxorubicin in cells was investigated using fluorescence microscopy with doxorubicin (red) and wheat germ agglutinin (WGA) membrane staining (green). Free doxorubicin treated groups display no or minimal doxorubicin uptake into cells below 2 µM in acidified pH media. We find that genipin cross-linked chitosan hydrogels exhibit fluorescence at the Ex/Em 570/650 nm used to visualize doxorubicin.^55,68^ Confocal images show that chitosan microparticles degraded from the hydrogel settle and collect around cells during the 48 h incubation, as confirmed by WGA staining and red fluorescence in doxorubicin negative control hydrogels (−DOX, 100 mM NaHCO_3_). Images of cell exposure to the control hydrogel without doxorubicin, containing only sodium bicarbonate (−DOX, 100 mM NaHCO_3_) do not show particle internalization of chitosan, confirming doxorubicin as a source of fluorescence in the drug-loaded trials. Cells exposed to dual-loaded doxorubicin and sodium bicarbonate hydrogels display increased co-localization of fluorescence in the cells (MDA-MB-231) and in the nuclei (MCF-7). The hydrogel releases bicarbonate ions alongside doxorubicin to neutralize the extracellular acid and prevent ion trapping. The free doxorubicin cases are absent of the buffer, resulting in acids inhibiting doxorubicin cell uptake. Therefore, the hydrogel cases achieve a more efficient doxorubicin diffusion into the cell due to the neutralization of the pH. Comparing doxorubicin accumulation with hydrogels to free doxorubicin uptake in cells from 0 to 5 µM at low pH confirms the feasibility of increasing doxorubicin efficacy with dual loading of sodium bicarbonate and doxorubicin. These results provide evidence that the dual-loaded hydrogels are more effective at increasing cellular uptake of doxorubicin than doxorubicin alone by providing an alkaline environment during doxorubicin delivery, preventing pH-dependent chemoresistance. This can also overcome the limitations of high dosage requirements of doxorubicin under an acidic extracellular environment, reducing the toxicity of doxorubicin to local healthy tissues.

**Figure 8:**
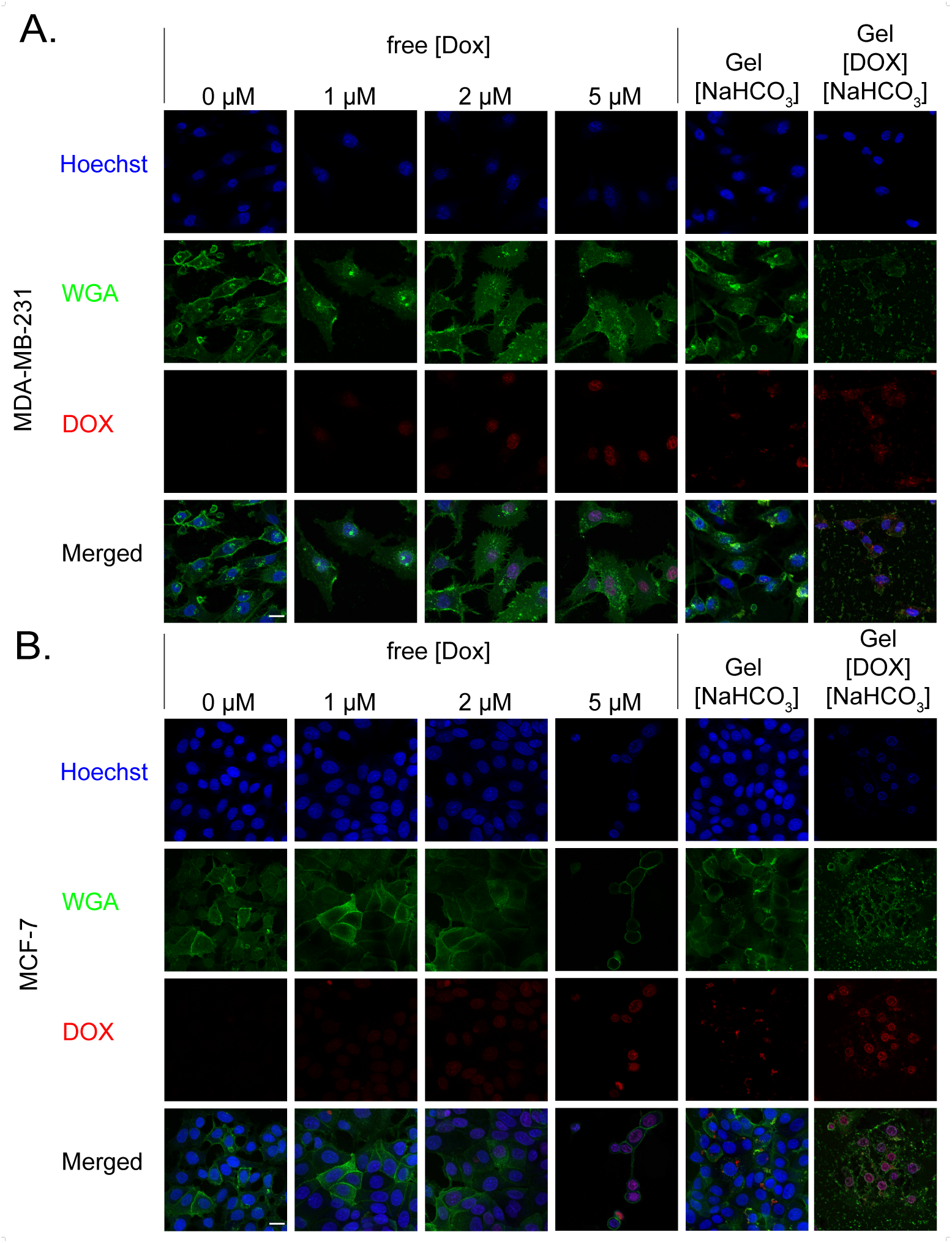
Cell internalization of free doxorubicin (0-5 µM) and doxorubicin released from sodium bicarbonate loaded chitosan-PEG hydrogels at low pH (6.5) at 48 h. The pH regulating hydrogels were fabricated with a sodium bicarbonate concentration of 100 mM and doxorubicin concentration of 50 µM. This gives a doxorubicin concentration of 0.47 *±* 0.05 µM in the cell solution over a 48 h period. Non-treated cells and cells treated with doxorubicin-free hydrogels (100 mM sodium bicarbonate) were used as controls. (A-B) Fluorescence microscopy images of MDA-MB-231 and MCF-7 cells show the cellular uptake of doxorubicin from pH regulating hydrogels is higher than cellular uptake of free doxorubicin. The nuclei were stained with Hoechst, Alexa 488-conjugated WGA was used to label the cell membrane, and (last column) overlay of all staining. All cells were imaged using a confocal fluorescence microscope (Olympus) with a 20x objective. Scale bar in bottom left image: 20 µm.

## Conclusion

We have presented a biocompatible, biodegradable, non-toxic, physical and covalent cross-linked chitosan-PEG based hydrogel system that can encapsulate and deliver sodium bicarbonate and doxorubicin. Sodium bicarbonate released from the hydrogel increases the local environment pH to a physiological level, causing an increase in chemotherapeutic efficacy in our in vitro experiments. We have conducted experiments on MDA-MB-231 and MCF-7 breast cancer cells to measure cancer cell viability and doxorubicin uptake to show that the doxorubicin and sodium bicarbonate releasing hydrogels have a potential to significantly improve chemotherapeutic treatment efficacy. The hydrogel in this work is injectable and can be locally delivered through injection. The shelf-life experiments show that the hydrogel components can be stored in a refrigerator for the long-term without affecting its mechanical and release properties.

We acknowledge that our study is in vitro, and for clinical applications, in vivo trials are expected. Nonetheless, our in vitro work, data and findings presented in the paper should help design future in vivo trials. The pH-regulating injectable hydrogel platforms such as the one presented here can potentially improve the uptake of other weak base chemotherapeutics that are susceptible to ion trapping, reducing harmful side effects from high dosage. In addition, due to rising evidence of the negative impact of acidosis on immune evasion, similar pH regulating hydrogel platforms can be used to improve immunotherapy outcomes.^69–73^

## Experimental Section

### Preparation of hydrogels

Chitosan hydrogels were fabricated based on protocols adapted from.^48,55,74,75^ Chitosan (MilliporeSigma, 2.5% wt/vol) was dissolved in glacial acetic acid solution (MilliporeSigma, 1% vol/vol) by magnetic stirring for 24 h. After dissolving, the pH of the chitosan solution was adjusted, and it was then sterilized by autoclaving (122°C, 15 min). PEG (MilliporeSigma, 75% wt/vol in deionized water) and genipin (Ambeed Inc, 7.5% wt/vol in 100% ethanol) were sterilized using 0.22 µm pore syringe filters. A mixture of chitosan (1.5% wt/vol), PEG (5% wt/vol), genipin (0.1% wt/vol), doxorubicin (Selleck Chemicals), and deionized water were vortex mixed in conical tubes and cooled at 4°C for 1 h. Next, the solutions were poured into petri dishes after the addition of sodium bicarbonate (MilliporeSigma) at indicated concentrations. The dishes were incubated at 37°C for 24 h to facilitate the gelation. Finally, 8 mm diameter, 100 mm^3^ volume biopsy punches were taken from the hydrogels and triple washed with PBS (MilliporeSigma) for experiments.

### Fourier-transform infrared spectroscopy

FTIR scans were obtained using a Spectrum Two FT-IR Spectrometer (PerkinElmer) to investigate the structural changes induced in the hydrogel by cross-linking. Synthesized hydrogels were frozen at −80 °C for 4 h before placement in a freeze-dryer (Labconco) with the condenser set at −50 °C. The hydrogels were allowed to dry for 48 h before analysis using the Universal ATR accessory (UATR, PerkinElmer). FTIR spectrum of the hydrogel constituents were obtained by mixing 2 mg of constituents with 98 mg of KBr. The 100 mg mixture was mashed before placing in an Evacuable Pellet Die (Specac) and pressed using a manual hydraulic press (Specac) at a force of 10 ton (US). FTIR spectra in the wavenumber range from 4,000 to 400 cm*^−^*^1^ were recorded in triplicates. Menges Spectragryph software was then used to normalize the peaks and average spectrum across triplicates.

### Scanning electron microscopy

Hydrogel samples were immersed with a fixative consisting of 5% glutaraldehyde (Millipore-Sigma) in 0.1 M cacodyolate buffer (Electron Microscopy Sciences) at room temperature for 1 h. After fixation, the samples were washed three times for 20 min in 0.1 M cacodylate buffer and sectioned to expose the inner cross-section. The samples were then dehydrated through an ethanol (Fisher Scientific) series from 30% to 100% ethanol in increments of 5% with three consecutive ethanol changes at 100%. The samples were left in 100% ethanol for 24 h before critical point drying using CO_2_. Samples were mounted on aluminum specimen stubs using double-sided adhesive carbon tape (Electron Microscopy Sciences), and then a thin layer of gold was deposited onto the sample using a sputter coater (Emitech K550). Images were captured at 2.00 kV and a working distance of 10 mm with a Thermo Apreo VS SEM.

### Rheological characterization of hydrogels

Rheological experiments were performed on an ARES G2 Rheometer using a parallel plate geometry of 25 mm diameter. All experiments were performed at 37°C and 1% strain. Oscillatory shear experiments were evaluated over the frequency range of 1 to 100 rad/s. The viscosity was measured as a function of the shear strain rate from 0.1 to 100 rad/s. Time sweeps were conducted at 100 rad/s.

### Sodium bicarbonate release studies

The pH of PBS and cell media samples were measured using the Mettler Toledo FiveEasy pH meter and electrode. To evaluate bicarbonate release, hydrogels were placed in 24-well plates (Celltreat) containing 2 mL of 1×PBS release media adjusted to pH 6. The plates were stored at 37 °C, and the pH was measured at predetermined time intervals.

### Doxorubicin release studies

For release studies, doxorubicin and bicarbonate loaded hydrogels were placed in well plates containing 2 mL 1× PBS release media adjusted to pH (7.4, 6.0 and 4.5). All the experiments were carried out in triplicate. The well plates were stored at 37 °C and sampled at predetermined time intervals. 200 µL samples were withdrawn, and the volume was re-placed with 200 µL of fresh PBS at the indicated pH. Doxorubicin fluorescence (480 nm excitation, 590 nm emission peak) was quantified with the Cytation 5 manager plate reader (BioTek^®^ Instruments, Inc., Winooski, VT, USA). The doxorubicin standard curve was prepared by quantifying the fluorescence of a serial dilution in PBS from 1.0 *×* 10^3^ to 1.2 *×* 10*^−^*^4^ µM. The initial doxorubicin loading concentration was based on fluorescence detection of uncross-linked hydrogel mixtures.

### Swelling tests

The swelling ratios of hydrogels were studied by weighing hydrogel sample punch outs before and after swelling in PBS (pH 7.4, 6.0 and 4.5) for 24 h at 37 °C. The hydrogels at post-swelling equilibrium were blotted to remove excess solution from the surface prior to weighing. The pre-swelled hydrogel and the swollen hydrogel weights of the samples were used to calculate the swelling ratio.

### Shelf-storage studies

The effect of storage on hydrogel samples was assessed by storing the combined hydrogel constituents within syringes followed by experimentation. The storage took place in either a refrigerator (4 °C) or a freezer (−20 °C). In the case of storage in the 4 °C refrigerator, the cross-linkers genipin and sodium bicarbonate were stored separately from the other hydrogel constituents. For storage in the −20 °C freezer, all constituents can be mixed in a single syringe and then stored frozen and thawed before use. The day of initial hydrogel preparation was designated as day 0 of storage. Prior to conducting doxorubicin release studies and rheological characterizations, the hydrogel underwent a 24 h cross-linking period at 37 °C in petri dishes.

### Cell culture

Human breast cancer cell lines MDA-MB-231 (ATCC) and MCF-7 (ATCC) were grown in DMEM/F12 (Gibco) with 15 mM HEPES supplemented with 10% v/v fetal bovine serum (VWR) and 1% v/v Penicillin-Streptomycin solution (Gibco). Human breast epithelial cell line MCF-10A (ATCC) was grown in complete DMEM/F12 (Gibco) supplemented with 5% v/v horse serum (Gibco), 1% v/v Penicillin-Streptomycin solution (Gibco), 20 ng/ml EGF (Peprotech), 0.5 mg/ml hydrocortisone (MilliporeSigma), 10 µg/ml Insulin (MilliporeSigma) and 100 ng/mL cholera toxin (MilliporeSigma). Cells were expanded in T-25 or T-75 flasks (Cell star, USA) at 37°C in a humidified incubator containing 5% CO_2_. Cell media was changed every 2 days and cells were passaged at 80% confluence. All experiments were performed with cells between passages 4 and 10.

### Hydrogel toxicity test

MCF-10A cells were seeded in 24-well plates at a density of 100,000 cells/well and incubated at 37°C for 24 hours in 2 mL complete DMEM/F12/well to produce adherent monolayers. Media was exchanged for 2% serum DMEM/F12, and hydrogel samples were rinsed with PBS and carefully placed into 12-well plate inserts (VWR). Well inserts were then placed into wells of adherent cell cultures. Well plates were incubated for an additional 24 h after hydrogel exposure. Following incubation, gentle aspiration was performed to remove media and hydrogels from the wells, and the viability of remaining cells was quantified via MTT assay. Cells were incubated in serum and phenol-red-free media with MTT (Gibco) working solution (0.5 mg/ml) for 3 h. After MTT crystallization, the working solution was aspirated and 100% Dimethylsulfoxide (DMSO, MilliporeSigma) was used to dissolve the formazan crystals. Plates were incubated at room temperature for 15 mins in the dark. Absorbance was measured at 570 nm and a reference wavelength of 650 nm. Cell viability was calculated as the noise-corrected reading (*Ab*_570_ *− Ab*_650_) for each sample normalized to the noise-corrected reading of the control case.

### pH dependent IC**_50_** assay

MDA-MB-231 and MCF-7 cells were cultured in 12-well clear, flat-bottom microplates (Cell-Treat), at a density of 1×10^6^ cells per well in 2 mL of complete culture medium for 24 h. Titrated amounts of doxorubicin (max concentration 10 µM) in pH 7.4 and pH 6.5 2% serum media was then added to the cells. After 24 h, an MTT assay was performed to evaluate cell metabolic activity (as a measure of cell viability). The drug medium was carefully re-placed by fresh serum and phenol-red-free media and 200 µL MTT reagent (0.5 mg/ml) was added. The MTT reagent was prepared in sterile 1× PBS at 5 mg/ml. After 3 h of incubation at 37°C, the well plates were centrifuged, the MTT solution was gently aspirated, and DMSO was added to dissolve the formazan crystals. The absorbance was measured at a wavelength of 570 nm and a reference wavelength of 650 nm. The half-maximal inhibitory concentration (IC_50_) was determined for each pH by fitting the data to a four-parameter dose-response model in GraphPad Prism (version 9.5.1). This process was repeated using bicarbonate-loaded hydrogels with pH 6.5 2% serum media and MTT assay performed after 48 h.

### Doxorubicin uptake studies

To study the uptake of doxorubicin into cells, MDA-MB-231 cells and MCF-7 cells were transferred at a density of 1×10^5^ cells/mL to fibronectin (MilliporeSigma, 2.5 µg/cm^2^) coated glass bottom petri dishes and allowed to adhere for 24 h. The cell culture media was replaced with 2 mL 2% serum media adjusted to pH 6.5 for the low pH cases. Sodium bicarbonate containing hydrogels (200 mM) were added to the same media for the high pH cases. A 0.5 to 2 µM range of concentrations of the intrinsic fluorescent doxorubicin was applied to cells for 48 h. Cell monolayers were then rinsed three times with PBS and fixed with 4% paraformaldehyde (VWR) for 15 min. Next, the samples were rinsed three times with PBS before counter staining with Hoechst 33342 (Thermo Fisher Scientific) overnight at 4°C. Fluorescent images were captured by an Olympus FV3000 Confocal Microscope in both blue (Excitation/Emission, 405/470 nm) and red (Excitation/Emission, 480/590 nm) channels. The fluorescence intensity of doxorubicin in the nucleus was quantified via the two channels in ImageJ. First, thresholds were applied to the images to define the borders of the cell nucleus in the blue channel. Then the integrated density of doxorubicin in the red channel was calculated according to the nucleus region. The corrected total doxorubicin cell fluorescence in the nucleus was calculated by subtracting background fluorescence.

### Doxorubicin uptake with doxorubicin and sodium bicarbonate loaded hydrogels

MDA-MB-231 and MCF-7 cells were passaged and resuspended at a density of 1×10^5^ cells/mL. 750 µL of cell suspension was added into the well of a fibronectin coated glass bottom petri dish and incubated overnight. Cell culture media was replaced with 2 mL 2% serum media adjusted to pH 6.5. Prior to co-incubation with cells, the hydrogels were then rinsed three times with PBS and placed in cell culture well inserts (MilliporeSigma). After 48 h of treatment, the hydrogels were removed and cells were washed three times with PBS.

Cell monolayers were fixed with 4% paraformaldehyde (VWR) for 15 min, washed three times with PBS for 5 min, stained with Alexa Fluor 488 wheat germ agglutinin (WGA, Thermo Fisher Scientific) at 5 µg/mL for 10 mins, and rinsed twice with PBS. Samples were then stained overnight with Hoechst 33342 at 5 µg/mL at 4°C and imaged using a confocal microscope with a 20x objective magnification.

### Statistical analysis

All analysis were performed in Prism 9 (GraphPad) and data are reported as mean ± standard error of the mean. Individual comparisons were made using an unpaired t-test. One way analysis of variance (ANOVA) followed by Tukey’s HSD post hoc test was used for data sets with multiple comparisons. A value of p *<* 0.05 was considered a statistically significant difference.

## Acknowledgement

This work was supported by Brown University School of Engineering and the Brown University Office of the Vice President of Research SEED Award to Dr. V. Srivastava. The authors thank Savan Santoki for the early set-up of hydrogel experiments, Dr. Roberto Zenit at Brown University for providing access to the ARES G2 Rheometer, Dr. Edith Mathiowitz at Brown University for providing access to FTIR analysis, and Geoffrey Williams from the Brown University Leduc Bioimaging Facility for providing help with microscopic imaging.

---

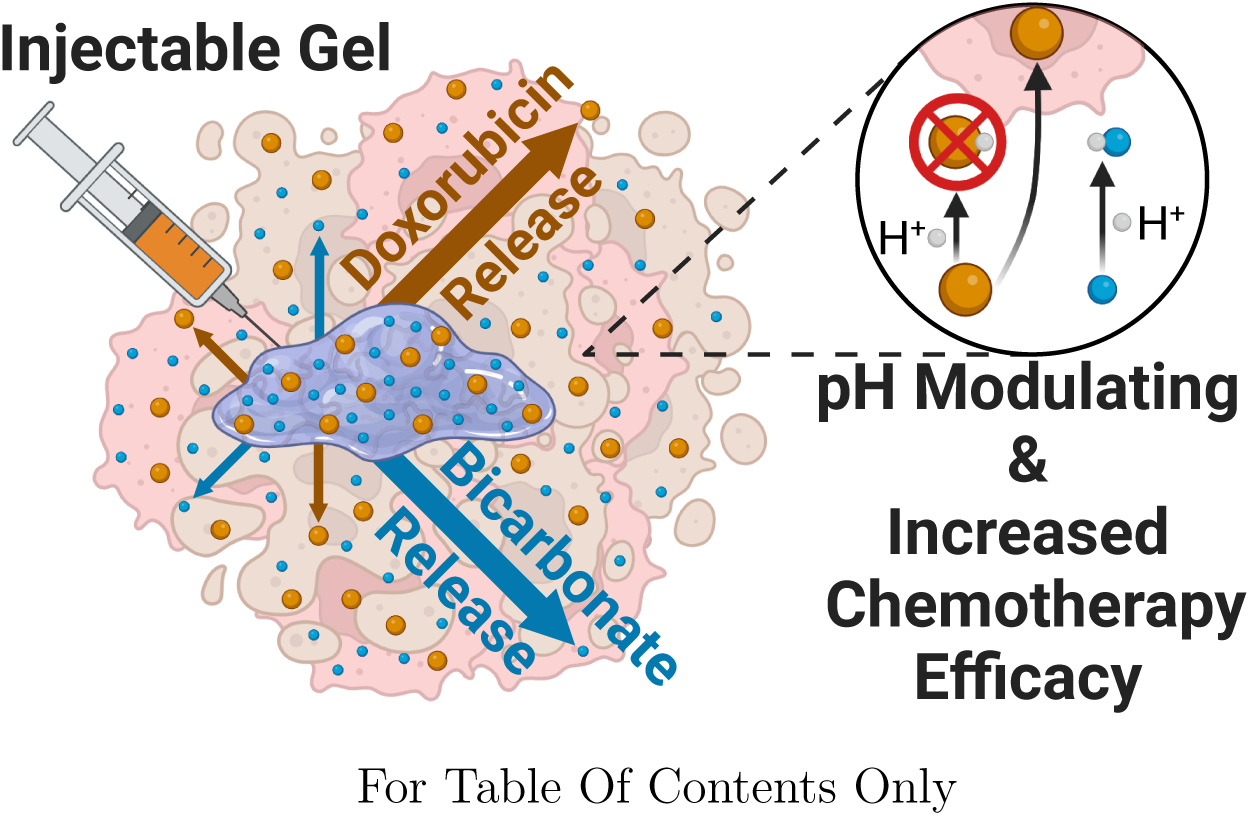

## Notes

### Competing Interest Statement

The authors have declared no competing interest.

### Summary of Updates

Abstract updated; Scheme 1 included; Results and Discussion updated; Figures 2 and 3 revised; Figure 4 included; Conclusion updated; Experimental Section updated.

## References

(1) Swietach, P.; Vaughan-Jones, R. D.; Harris, A. L.; Hulikova, A. The Chemistry, Physiology and Pathology of pH in Cancer. Philosophical Transactions of the Royal Society B: Biological Sciences 2014, 369, 20130099.

(2) Damaghi, M.; Wojtkowiak, J. W.; Gillies, R. J. pH Sensing and Regulation in Cancer. Frontiers in Physiology 2013, 4, 370.

(3) Persi, E.; Duran-Frigola, M.; Damaghi, M.; Roush, W. R.; Aloy, P.; Cleveland, J. L.; Gillies, R. J.; Ruppin, E. Systems Analysis of Intracellular pH Vulnerabilities for Cancer Therapy. Nature Communications 2018, 9, 2997.

(4) Boedtkjer, E.; Pedersen, S. F. The Acidic Tumor Microenvironment as a Driver of Cancer. Annual Review of Physiology 2020, 82, 103–126.

(5) Pillai, S. R.; Damaghi, M.; Marunaka, Y.; Spugnini, E. P.; Fais, S.; Gillies, R. J. Causes, Consequences, and Therapy of Tumors Acidosis. Cancer and Metastasis Reviews 2019, 38, 205–222.

(6) Luengo, A.; Li, Z.; Gui, D. Y.; Sullivan, L. B.; Zagorulya, M.; Do, B. T.; Ferreira, R.; Naamati, A.; Ali, A.; Lewis, C. A.; Thomas, C. J.; Spranger, S.; Matheson, N. J.; Vander Heiden, M. G. Increased Demand for NAD+ Relative to ATP Drives Aerobic Glycolysis. Molecular Cell 2021, 81, 691–707.

(7) Liberti, M. V.; Locasale, J. W. The Warburg Effect: How Does it Benefit Cancer Cells? Trends in Biochemical Sciences 2016, 41, 211–218.

(8) Fadaka, A.; Ajiboye, B.; Ojo, O.; Adewale, O.; Olayide, I.; Emuowhochere, R. Biology of Glucose Metabolization in Cancer Cells. Journal of Oncological Sciences 2017, 3, 45–51.

(9) Boedtkjer, E. Na+,HCO3-Cotransporter NBCn1 Accelerates Breast Carcinogenesis. Cancer and Metastasis Reviews 2019, 38, 165–178.

(10) Ferrari, S.; Perut, F.; Fagioli, F.; Brach Del Prever, A.; Meazza, C.; Parafioriti, A.; Picci, P.; Gambarotti, M.; Avnet, S.; Baldini, N.; Fais, S. Proton Pump Inhibitor Chemosensitization in Human Osteosarcoma: From the Bench to the Patients’ Bed. Journal of Translational Medicine 2013, 11, 268.

(11) Lee, S.; Axelsen, T. V.; Andersen, A. P.; Vahl, P.; Pedersen, S. F.; Boedtkjer, E. Disrupting Na+,HCO3–Cotransporter NBCn1 (Slc4a7) Delays Murine Breast Cancer Development. Oncogene 2016, 35, 2112–2122.

(12) Czowski, B. J.; Romero-Moreno, R.; Trull, K. J.; White, K. A. Cancer and pH Dynamics: Transcriptional Regulation, Proteostasis, and the Need for New Molecular Tools. Cancers 2020, 12, 2760.

(13) Krieg, B. J.; Taghavi, S. M.; Amidon, G. L.; Amidon, G. E. In Vivo Predictive Dissolution: Transport Analysis of the CO_2_, Bicarbonate In Vivo Buffer System. Journal of Pharmaceutical Sciences 2014, 103, 3473–3490.

(14) Parks, S. K.; Chiche, J.; Pouysśegur, J. Disrupting Proton Dynamics and Energy Metabolism for Cancer Therapy. Nature Reviews Cancer 2013, 13, 611–623.

(15) Swietach, P. What is pH Regulation, and Why Do Cancer Cells Need It? Cancer and Metastasis Reviews 2019, 38, 5–15.

(16) Chiche, J.; Ilc, K.; Laferrìere, J.; Trottier, E.; Dayan, F.; Mazure, N. M.; Brahimi-Horn, M. C.; Pouysśegur, J. Hypoxia-Inducible Carbonic Anhydrase IX and XII Promote Tumor Cell Growth by Counteracting Acidosis through the Regulation of the Intracellular pH. Cancer Research 2009, 69, 358–368.

(17) Rohani, N.; Hao, L.; Alexis, M. S.; Joughin, B. A.; Krismer, K.; Moufarrej, M. N.; Soltis, A. R.; Lauffenburger, D. A.; Yaffe, M. B.; Burge, C. B.; Bhatia, S. N.; Gertler, F. B. Acidification of Tumor at Stromal Boundaries Drives Transcriptome Alterations Associated with Aggressive Phenotypes. Cancer Research 2019, 79, 1952– 1966.

(18) Tavares-Valente, D.; Sousa, B.; Schmitt, F.; Baltazar, F.; Queiŕos, O. Disruption of pH Dynamics Suppresses Proliferation and Potentiates Doxorubicin Cytotoxicity in Breast Cancer Cells. Pharmaceutics 2021, 13, 242.

(19) Thews, O.; Riemann, A. Tumor pH and Metastasis: A Malignant Process Beyond Hypoxia. Cancer and Metastasis Reviews 2019, 38, 113–129.

(20) Senthebane, D. A.; Rowe, A.; Thomford, N. E.; Shipanga, H.; Munro, D.; Al Mazeedi, M. A.; Almazyadi, H. A.; Kallmeyer, K.; Dandara, C.; Pepper, M. S.; Parker, M. I.; Dzobo, K. The Role of Tumor Microenvironment in Chemoresistance: To Survive, Keep Your Enemies Closer. International Journal of Molecular Sciences 2017, 18, 1586.

(21) Huber, V.; Camisaschi, C.; Berzi, A.; Ferro, S.; Lugini, L.; Triulzi, T.; Tuccitto, A.; Tagliabue, E.; Castelli, C.; Rivoltini, L. Cancer Acidity: An Ultimate Frontier of Tumor Immune Escape and a Novel Target of Immunomodulation. Seminars in Cancer Biology 2017, 43, 74–89.

(22) Li, S.; Xiong, N.; Peng, Y.; Tang, K.; Bai, H.; Lv, X.; Jiang, Y.; Qin, X.; Yang, H.; Wu, C.; Zhou, P.; Liu, Y. Acidic pHe Regulates Cytoskeletal Dynamics through Conformational Integrin *β*1 Activation and Promotes Membrane Protrusion. Biochimica et Biophysica Acta (BBA) - Molecular Basis of Disease 2018, 1864, 2395–2408.

(23) Bogdanov, A.; Bogdanov, A.; Chubenko, V.; Volkov, N.; Moiseenko, F.; Moiseyenko, V. Tumor Acidity: From Hallmark of Cancer to Target of Treatment. Frontiers in Oncology 2022, 12, 979154.

(24) Raghunand, N.; He, X.; van Sluis, R.; Mahoney, B.; Baggett, B.; Taylor, C. W.; Paine-Murrieta, G.; Roe, D.; Bhujwalla, Z. M.; Gillies, R. J. Enhancement of Chemotherapy by Manipulation of Tumour pH. British Journal of Cancer 1999, 80, 1005–1011.

(25) Trebinska-Stryjewska, A.; Swiech, O.; Opuchlik, L. J.; Grzybowska, E. A.; Bilewicz, R. Impact of Medium pH on DOX Toxicity Toward HeLa and A498 Cell Lines. ACS Omega 2020, 5, 7979–7986.

(26) Gu, Y.; Zhao, Z.; Niu, G.; Zhang, H.; Wang, Y.; Kwok, R. T.; Lam, J. W.; Tang, B. Z. Visualizing Semipermeability of the Cell Membrane Using a pH-Responsive Ratiometric AIEgen. Chemical Science 2020, 11, 5753–5758.

(27) Pilon-Thomas, S.; Kodumudi, K. N.; El-Kenawi, A. E.; Russell, S.; Weber, A. M.; Luddy, K.; Damaghi, M.; Wojtkowiak, J. W.; Muĺe, J. J.; Ibrahim-Hashim, A.; Gillies, R. J. Neutralization of Tumor Acidity Improves Antitumor Responses to Immunotherapy. Cancer Research 2016, 76, 1381–1390.

(28) Robey, I. F.; Baggett, B. K.; Kirkpatrick, N. D.; Roe, D. J.; Dosescu, J.; Sloane, B. F.; Hashim, A. I.; Morse, D. L.; Raghunand, N.; Gatenby, R. A.; Gillies, R. J. Bicarbonate Increases Tumor pH and Inhibits Spontaneous Metastases. Cancer Research 2009, 69, 2260.

(29) Robey, I. F.; Martin, N. K. Bicarbonate and Dichloroacetate: Evaluating pH Altering Therapies in a Mouse Model for Metastatic Breast Cancer. BMC Cancer 2011, 11, 235.

(30) Ibrahim-Hashim, A.; Cornnell, H. H.; Abrahams, D.; Lloyd, M.; Bui, M.; Gillies, R. J.; Gatenby, R. A. Systemic Buffers Inhibit Carcinogenesis in TRAMP Mice. Journal of Urology 2012, 188, 624–631.

(31) Faes, S.; Dormond, O. Systemic Buffers in Cancer Therapy: The Example of Sodium Bicarbonate; Stupid Idea or Wise Remedy? Medicinal Chemistry 2015, 5, 540–544.

(32) Martin, N. K.; Robey, I. F.; Gaffney, E. A.; Gillies, R. J.; Gatenby, R. A.; Maini, P. K. Predicting the Safety and Efficacy of Buffer Therapy to Raise Tumour pHe: An Integrative Modelling Study. British Journal of Cancer 2012, 106, 1280–1287.

(33) Gillies, R. J.; Pilot, C.; Mahipal, A. Buffer Therapy → Buffer Diet. Journal of Nutrition & Food Sciences 2018, 8, 1000685.

(34) de Lourdes Ribeiro, M. C.; Silva, A. S.; Bailey, K. M.; Kumar, N. B.; Sellers, T. A.; Gatenby, R. A.; Ibrahim-Hashim, A.; Gillies, R. J.; Author, C.; Member, S.; Lee, H. Buffer Therapy for Cancer. Journal of Nutrition & Food Sciences 2012, 2, 6.

(35) Abumanhal-Masarweh, H.; Koren, L.; Zinger, A.; Yaari, Z.; Krinsky, N.; Kaneti, G.; Dahan, N.; Lupu-Haber, Y.; Suss-Toby, E.; Weiss-Messer, E.; Schlesinger-Laufer, M.; Shainsky-Roitman, J.; Schroeder, A. Sodium Bicarbonate Nanoparticles Modulate the Tumor pH and Enhance the Cellular Uptake of Doxorubicin. Journal of Controlled Release 2019, 296, 1–13.

(36) Lam, S. F.; Bishop, K. W.; Mintz, R.; Fang, L.; Achilefu, S. Calcium Carbonate Nanoparticles Stimulate Cancer Cell Reprogramming to Suppress Tumor Growth and Invasion in an Organ-on-a-Chip System. Scientific Reports 2021, 11, 9246.

(37) Riganti, C.; Gazzano, E.; Gulino, G. R.; Volante, M.; Ghigo, D.; Kopecka, J. Two Repeated Low Doses of Doxorubicin are More Effective than a Single High Dose Against Tumors Overexpressing P-glycoprotein. Cancer Letters 2015, 360, 219–226.

(38) Pugazhendhi, A.; Edison, T. N. J. I.; Velmurugan, B. K.; Jacob, J. A.; Karuppusamy, I. Toxicity of Doxorubicin (Dox) to Different Experimental Organ Systems. Life Sciences 2018, 200, 26–30.

(39) Eikenberry, S. A Tumor Cord Model for Doxorubicin Delivery and Dose Optimization in Solid Tumors. Theoretical Biology & Medical Modelling 2009, 6, 16.

(40) Dawidczyk, C. M.; Russell, L. M.; Hultz, M.; Searson, P. C. Tumor Accumulation of Liposomal Doxorubicin in Three Murine Models: Optimizing Delivery Efficiency. *Nanomedicine: Nanotechnology*, Biology, and Medicine 2017, 13, 1637.

(41) Robert Li, Y.; Traore, K.; Zhu, H. Novel Molecular Mechanisms of Doxorubicin Cardiotoxicity: Latest Leading-Edge Advances and Clinical Implications. Molecular and Cellular Biochemistry 2023, 1, 1–12.

(42) Chen, M. et al. Multifunctional Microspheres Dual-Loaded with Doxorubicin and Sodium Bicarbonate Nanoparticles to Introduce Synergistic Trimodal Interventional Therapy. ACS Applied Bio Materials 2021, 4, 3476–3489.

(43) Chen, M.; Guo, X.; Shen, L.; Ding, J.; Yu, J.; Chen, X.; Wu, F.; Tu, J.; Zhao, Z.; Nakajima, M.; Song, J.; Shu, G.; Ji, J. Monodisperse CaCO_3_-Loaded Gelatin Microspheres for Reversing Lactic Acid-Induced Chemotherapy Resistance During TACE Treatment. International Journal of Biological Macromolecules 2023, 231, 123160.

(44) Muzzarelli, R. A.; El Mehtedi, M.; Bottegoni, C.; Aquili, A.; Gigante, A. Genipin-Crosslinked Chitosan Gels and Scaffolds for Tissue Engineering and Regeneration of Cartilage and Bone. Marine Drugs 2015, 13, 7314.

(45) Tang, G.; Tan, Z.; Zeng, W.; Wang, X.; Shi, C.; Liu, Y.; He, H.; Chen, R.; Ye, X. Recent Advances of Chitosan-Based Injectable Hydrogels for Bone and Dental Tissue Regeneration. Frontiers in Bioengineering and Biotechnology 2020, 8, 1084.

(46) Vo, N. T.; Huang, L.; Lemos, H.; Mellor, A. L.; Novakovic, K. Genipin-Crosslinked Chitosan Hydrogels: Preliminary Evaluation of the In Vitro Biocompatibility and Biodegradation. Journal of Applied Polymer Science 2021, 138, 50848.

(47) Klein, M. P.; Hackenhaar, C. R.; Lorenzoni, A. S.; Rodrigues, R. C.; Costa, T. M.; Ninow, J. L.; Hertz, P. F. Chitosan Crosslinked with Genipin as Support Matrix for Application in Food Process: Support Characterization and *β*-d-galactosidase Immobilization. Carbohydrate Polymers 2016, 137, 184–190.

(48) Liu, L.; Tang, X.; Wang, Y.; Guo, S. Smart Gelation of Chitosan Solution in the Presence of NaHCO_3_ for injectable drug delivery system. International Journal of Pharmaceutics 2011, 414, 6–15.

(49) Wahba, M. I. Sodium Bicarbonate-Gelled Chitosan Beads as Mechanically Stable Carriers for the Covalent Immobilization of Enzymes. Biotechnology Progress 2018, 34, 347–361.

(50) Ernesto, J. V.; Gasparini, Í d. M.; Corazza, F. G.; Mathor, M. B.; da Silva, C. F.; Leite-Silva, V. R.; Andŕeo-Filho, N.; Lopes, P. S. Physical, Chemical, and Biological Characterization of Biodegradable Chitosan Dressing for Biomedical Applications: Could Sodium Bicarbonate Act as a Crosslinking Agent? Materials Chemistry and Physics 2023, 301, 127636.

(51) Fletes-Vargas, G.; Espinosa-Andrews, H.; Cervantes-Uc, J. M.; Limón-Rocha, I.; Luna-Bárcenas, G.; Vázquez-Lepe, M.; Morales-Herńandez, N.; Jiménez-Ávalos, J. A.; Mejía-Torres, D. G.; Ramos-Martínez, P.; Rodŕíguez-Rodŕíguez, R. Porous Chitosan Hydrogels Produced by Physical Crosslinking: Physicochemical, Structural, and Cytotoxic Properties. Polymers 2023, 15, 2203.

(52) Ortega-Ortiz, H.; Gutíerrez-Rodŕíguez, B.; Cadenas-Pliego, G.; Jimenez, L. I. Antibacterial Activity of Chitosan and the Interpolyelectrolyte Complexes of Poly(acrylic acid)-Chitosan. Brazilian Archives of Biology and Technology 2010, 53, 623–628.

(53) Queiroz, M. F.; Melo, K. R. T.; Sabry, D. A.; Sassaki, G. L.; Rocha, H. A. O. Does the Use of Chitosan Contribute to Oxalate Kidney Stone Formation? Marine Drugs 2015, 13, 141–158.

(54) Reay, S. L.; Jackson, E. L.; Ferreira, A. M.; Hilkens, C. M.; Novakovic, K. In Vitro Evaluation of the Biodegradability of Chitosan–Genipin Hydrogels. Materials Advances 2022, 3, 7946–7959.

(55) Vo, N. T.; Huang, L.; Lemos, H.; Mellor, A.; Novakovic, K. Poly(ethylene glycol)-Interpenetrated Genipin-Crosslinked Chitosan Hydrogels: Structure, pH Responsiveness, Gelation Kinetics, and Rheology. Journal of Applied Polymer Science 2020, 137, 49259.

(56) Michl, J.; Park, K. C.; Swietach, P. Evidence-Based Guidelines for Controlling pH in Mammalian Live-Cell Culture Systems. Communications Biology 2019, 2, 144.

(57) Mahoney, B. P.; Raghunand, N.; Baggett, B.; Gillies, R. J. Tumor Acidity, Ion Trapping and Chemotherapeutics: I. Acid pH Affects the Distribution of Chemotherapeutic Agents in Vitro. Biochemical Pharmacology 2003, 66, 1207–1218.

(58) Wojtkowiak, J. W.; Verduzco, D.; Schramm, K. J.; Gillies, R. J. Drug Resistance and Cellular Adaptation to Tumor Acidic pH Microenvironment. Molecular pharmaceutics 2011, 8, 2032.

(59) Wan, X.; Hou, J.; Liu, S.; Zhang, Y.; Li, W.; Zhang, Y.; Ding, Y. Estrogen Receptor *α* Mediates Doxorubicin Sensitivity in Breast Cancer Cells by Regulating E-Cadherin. Frontiers in Cell and Developmental Biology 2021, 9, 583572.

(60) Lovitt, C. J.; Shelper, T. B.; Avery, V. M. Doxorubicin Resistance in Breast Cancer Cells is Mediated by Extracellular Matrix Proteins. BMC Cancer 2018, 18, 41.

(61) Bao, L.; Haque, A.; Jackson, K.; Hazari, S.; Moroz, K.; Jetly, R.; Dash, S. Increased Expression of P-Glycoprotein Is Associated with Doxorubicin Chemoresistance in the Metastatic 4T1 Breast Cancer Model. The American Journal of Pathology 2011, 178, 838.

(62) Bao, L.; Hazari, S.; Mehra, S.; Kaushal, D.; Moroz, K.; Dash, S. Increased Expression of P-Glycoprotein and Doxorubicin Chemoresistance of Metastatic Breast Cancer Is Regulated by miR-298. The American Journal of Pathology 2012, 180, 2490.

(63) Mirzaei, S.; Abadi, A. J.; Gholami, M. H.; Hashemi, F.; Zabolian, A.; Hushmandi, K.; Zarrabi, A.; Entezari, M.; Aref, A. R.; Khan, H.; Ashrafizadeh, M.; Samarghandian, S. The Involvement of Epithelial-to-Mesenchymal Transition in Doxorubicin Resistance: Possible Molecular Targets. European Journal of Pharmacology 2021, 908, 174344.

(64) Rauschner, M.; Lange, L.; Hüsing, T.; Reime, S.; Nolze, A.; Maschek, M.; Thews, O.; Riemann, A. Impact of the Acidic Environment on Gene Expression and Functional Parameters of Tumors In Vitro and In Vivo. Journal of Experimental and Clinical Cancer Research 2021, 40, 1–14.

(65) Aroui, S.; Ram, N.; Appaix, F.; Ronjat, M.; Kenani, A.; Pirollet, F.; De Waard, M. Maurocalcine as a Non Toxic Drug Carrier Overcomes Doxorubicin Resistance in the Cancer Cell Line MDA-MB 231. Pharmaceutical Research 2009, 26, 836–845.

(66) Aroui, S.; Brahim, S.; Waard, M. D.; Kenani, A. Cytotoxicity, Intracellular Distribution and Uptake of Doxorubicin and Doxorubicin Coupled to Cell-Penetrating Peptides in Different Cell Lines: A Comparative Study. Biochemical and Biophysical Research Communications 2010, 391, 419–425.

(67) Condello, M.; Cosentino, D.; Corinti, S.; Di Felice, G.; Multari, G.; Gallo, F. R.; Arancia, G.; Meschini, S. Voacamine Modulates the Sensitivity to Doxorubicin of Resistant Osteosarcoma and Melanoma Cells and Does Not Induce Toxicity in Normal Fibroblasts. Journal of Natural Products 2014, 77, 855–862.

(68) Hoemann, C. D.; Guzmán-Morales, J.; Tran-Khanh, N.; Lavalĺee, G.; Jolicoeur, M.; Lavertu, M. Chitosan Rate of Uptake in HEK293 Cells is Influenced by Soluble versus Microparticle State and Enhanced by Serum-Induced Cell Metabolism and Lactate-Based Media Acidification. Molecules 2013, 18, 1015–1035.

(69) Huntington, K. E.; Louie, A.; Zhou, L.; Seyhan, A. A.; Maxwell, A. W.; El-Deiry, W. S. Colorectal Cancer Extracellular Acidosis Decreases Immune Cell Killing and is Partially Ameliorated by pH-Modulating Agents that Modify Tumor Cell Cytokine Profiles. American Journal of Cancer Research 2022, 12, 138–151.

(70) Kay, E. J.; Zanivan, S. Bicarbonate Transport as Regulator of Antitumour Immunity in Pancreatic Cancer. Molecular Oncology 2023, 17, 541–544.

(71) Yuan, Y. H.; Zhou, C. F.; Yuan, J.; Liu, L.; Guo, X. R.; Wang, X. L.; Ding, Y.; Wang, X. N.; Li, D. S.; Tu, H. J. NaHCO_3_ Enhances the Antitumor Activities of Cytokine-Induced Killer Cells Against Hepatocellular Carcinoma HepG2 Cells. Oncology Letters 2016, 12, 3167–3174.

(72) Damgaci, S.; Ibrahim-Hashim, A.; Enriquez-Navas, P. M.; Pilon-Thomas, S.; Guvenis, A.; Gillies, R. J. Hypoxia and Acidosis: Immune Suppressors and Therapeutic Targets. Immunology 2018, 154, 354–362.

(73) seung Jin, H.; som Choi, D.; Ko, M.; Kim, D.; hee Lee, D.; Lee, S.; Lee, A. Y.; Kang, S. G.; Kim, S. H.; Jung, Y.; Jeong, Y.; Chung, J. J.; Park, Y. Extracellular pH Modulating Injectable Gel for Enhancing Immune Checkpoint Inhibitor Therapy. Journal of Controlled Release 2019, 315, 65–75.

(74) Sharma, P. K.; Halder, M.; Srivastava, U.; Singh, Y. Antibacterial PEG-Chitosan Hydrogels for Controlled Antibiotic/Protein Delivery. ACS Applied Bio Materials 2019, 5313–5322.

(75) Heimbuck, A. M.; Priddy-Arrington, T. R.; Padgett, M. L.; Llamas, C. B.; Barnett, H. H.; Bunnell, B. A.; Caldorera-Moore, M. E. Development of Responsive Chitosan-Genipin Hydrogels for the Treatment of Wounds. ACS Applied Bio Materials 2019, 2, 2879–2888.

